# RNA modulation of transport properties and stability in phase separated condensates

**DOI:** 10.1101/2021.03.05.434111

**Authors:** Andrés R. Tejedor, Adiran Garaizar, Jorge Ramírez, Jorge R. Espinosa

**Author notes:** Electronic mail (Jorge Ramírez). Electronic mail.

## Abstract

One of the key mechanisms employed by cells to control their spatiotemporal organization is the formation and dissolution of phase-separated condensates. The balance between condensate assembly and disassembly can be critically regulated by the presence of RNA. In this work, we use a novel sequence-dependent coarse-grained model for proteins and RNA to unravel the impact of RNA in modulating the transport properties and stability of biomolecular condensates. We explore the phase behavior of several RNA-binding proteins such as FUS, hnRNPA1 and TDP-43 proteins along with that of their corresponding prion-like domains and RNA-recognition motifs, from absence to moderately high RNA concentration. By characterising the phase diagram, key molecular interactions, surface tension and transport properties of the condensates, we report a dual RNA-induced behavior: On the one hand, RNA enhances phase separation at low concentration as long as the RNA radius of gyration is comparable to that of the proteins, whilst at high concentration it inhibits the ability of proteins to self-assemble independently of its length. On the other hand, along with the stability modulation, the viscosity of the condensates can be considerably reduced at high RNA concentration as long as the length of the RNA chains is shorter than that of the proteins. Conversely, long RNA strands increase viscosity, even at high concentration, but barely modify protein self-diffusion, which mainly depends on RNA concentration and on its own effect on droplet density. On the whole, our work rationalizes the different routes by which RNA can regulate phase separation and condensate dynamics, as well as the subsequent aberrant rigidification implicated in the emergence of various neuropathologies and age-related diseases.

## I. INTRODUCTION

Liquid-liquid phase separation (LLPS) is one of the key processes employed by cells to control the spatiotemporal organization of their many components^1–6^. This phenomenon – displayed by a large variety of biomolecules such as multivalent proteins and nucleic acids^6–11^ – is involved in wide-ranging aspects of the cell function such as membraneless compartmentalisation ^6,12–16^, signaling^2,17^, genome silencing^18–20^, formation of super-enhancers^21^, helping cells to sense and react to environmental changes^22^, or buffering cellular noise^23^ among many others^24–27^. The spontaneous demixing of the cell components into different coexisting liquid compartments occurs both inside the cytoplasm (e.g., P granules^1^ and RNA granules/bodies^28,29^) and in the cell nucleus (e.g., Cajal bodies^30^, nucleoli^31^, nuclear speckles^32,33^ or heterochromatin domains^19,20^), and enables the coordinated control of thousands of simultaneous chemical reactions that are required to maintain biological activity^34^. Beyond these diverse functionalities, membraneless organelles have also been observed to exert mechanical forces to induce chromatin reorganization^35,36^ or to act as molecular sensors of intracellular and extracellular exchanges^22^. Still novel biological roles, such as the propensity of condensates to buffer protein concentrations against gene expression noise continue being discovered^23,37^.

The biomolecular building blocks behind LLPS are usually proteins with intrinsically disordered regions (IDRs) or proteins with globular domains connected by flexible linkers that can establish multiple homotypic or heterotypic interactions with cognate biomolecules (e.g. a different IDR, RNA or DNA) over their interactions with the solvent^9,11^. Several DNA and RNA-binding proteins such as FUS^38–40^, hnRNPA1^15,16^, TDP-43^41,42^, LAF-1^43^, G3BP1^44–46^ or HP1^19,20^ have been observed to undergo phase separation both *in vivo* and *in vitro*. These proteins, besides their intrinsically disordered regions, frequently present additional specific domains with high physico-chemical affinity for RNA (termed as RNA recognition motifs (RRMs))^47^ or DNA^48^. In particular, the intermolecular binding between IDRs and RNA (either via specific RNA-RRM interactions or non-selective electrostatic and *π*-*π* interactions) have been found to be critical in regulating LLPS^43,49–54^.

*In vitro* experimental evidences show how protein aggregation can be enhanced upon addition of RNA at low concentration, whilst inhibited at high concentration^50,55,56^. Such reentrant behavior is in agreement with the hypothesis that solid-like aggregates are more readily formed in the cytoplasm than in the cell nucleus where the abundance of RNA is higher^57^. Moreover, besides modulating the stability of the condensates, RNA can affect their kinetic properties. A viscosity reduction of LAF-1 droplets (a key protein in P granules formation) after addition of short RNA strands has been observed without significantly affecting droplet stability^43^. On the contrary, the inclusion of long RNA chains inside the condensates can also enhance notably their viscosity at certain given concentrations^49,58^. Such RNA-induced modulation of droplet viscoelasticity (and recently observed by DNA^59^) is crucial in the regulation/dysregulation of the liquid-like behavior of RNA-binding proteins (RBPs) such as FUS^38–40^, hnRNPA1^15,16^, TDP-43^41,42,60^, TAF-15^57,61^ or EWSR1 among many others^51,57,61^. The resulting rigidification of these condensates can lead to the formation of pathological solid aggregates which are behind the onset of several neurodegenerative diseases such as amyotrophic lateral sclerosis (ALS), frontotemporal dementia or Alzheimer^15,62–66^. Because of that, a huge effort in understanding the underlying molecular factors involved in RNA-induced regulation of condensates stability and viscoelasticity is being devoted^8,12,53,54,67,68^.

Recent experimental advances in single-molecule Forster resonance energy transfer (smFRET) have enabled the direct observation of the structural and dynamic protein behavior in diluted conditions^69–71^, however, the thermodynamic and kinetic aspects inside the condensates are still hardly accessible^72,73^. Notably, particle tracking microrheology techniques have been successfully used to provide data about the mean squared displacement (MSD) of marked beads inside droplets, and then, via that MSD, condensate viscosity has been estimated^43,49,58,74^. Nevertheless, other fundamental magnitudes such as the protein mean squared displacement, end-to-end distance relaxation times, protein radius of gyration or droplet surface tensions are extremely challenging to obtain^53^. Moreover, direct measurements of the molecular contacts that promote phase separation are of great relevance, and rarely, this information can be unequivocally extracted^39,61,75^. The mutation and/or phosphorylation of specific residues along sequences can help in deciphering which contacts are key in sustaining LLPS^76,77^, but still a higher level of mechanistic and molecular resolution is needed.

In that respect, computer simulations emerge as a great tool to enlighten such blind spot^78–80^. The most recent advances in computational science have allowed to carry out impressive calculations mimicking *in vivo* conditions^81^. Atomistic Molecular Dynamics (MD) simulations have been also successfully proved in characterizing the conformational ensemble of single proteins and protein complexes^80,82,83^, pinpointing the link between chemical modifications and the modulation of protein–protein and protein-DNA interactions^84,85^, or guiding the development of chemically accurate coarse-grained models for LLPS^86–89^. Simultaneously, a huge effort in developing different levels of coarse-grained (CG) potentials, these including mean field models^90–93^, lattice-based simulations^94–97^, minimal models^98–102^, and sequence-dependent models^86,103,104^ is being devoted. By retaining the specific physico-chemical features of proteins, DNA and RNA, while averaging out others for computational efficiency, CG models have been widely used to elucidate key factors behind LLPS and their dependency on protein length^105,106^, amino acid sequence^86,103,107,108^, multivalency^94,109–114^, conformational flexibility^115,116^ and multicomponent composition^117–120^. Nevertheless, further work is needed regarding the role of RNA in LLPS^121^. On the one hand, atomistic MD simulations have provided binding free energies of specific protein/RNA complexes, but are limited to very few protein replicas^122,123^. On the other hand, coarse-grained models have been recently proposed to elucidate the effect of RNA on phase separation of small prion-like domains such as those of FUS^124^, protamine^125^ or LAF-1^119^. Remarkably, the work by Regy *et al.*^119^ presents a detailed parametrization of a CG model for RNA within the framework of the HPS protein potential^86^, opening up new possibilities to link the molecular mechanisms of RNA-RBP condensates to their macroscopic phase behavior.

The present work aims to narrow down this problem by shedding light on the RNA modulation of transport properties and stability of RBP condensates. By employing a high-resolution CG model for RNA and IDPs^86,119,126^, we explore the phase behaviour of different RNA-binding proteins which undergo LLPS such as FUS, hnRNPA1 or TDP-43 as well as their corresponding prion-like domains and RNA-recognition motifs in absence *vs.* presence of poly-Uridine (poly-U) RNA. After validating the model against experimental saturation concentration trends of these pure proteins at physiological salt concentration, we characterize how RNA regulates the coexistence line of these condensates as a function of RNA concentration for a constant poly-U length, as well as for different strand lengths at a constant poly-U/protein concentration. Beyond evidencing RNA-induced reentrant phase separation^50,55–57^, we find a critical minimum length below which RNA cannot promote LLPS even at low concentration. Moreover, we characterize the transport properties (i.e., protein mobility and viscosity) of the condensates as a function of RNA saturation and length. While protein diffusion is predominantly controlled by RNA concentration rather than by strand length, the viscosity of the droplets is critically regulated by both factors, being RNA length a key element in LLPS. Taken together, our work provides a framework to rationalise from a molecular and thermodynamic perspective the ubiquitous dual effect of RNA in the stability and kinetics of RNA-RBP condensates.

## II. RESULTS

### A. Methods and sequence-dependent model validation

Biomolecular condensates are stabilized by chemically diverse weak protein–protein interactions, which are determined by the specific nature (e.g., hydrophobicity, aromaticity, and charge) of the encoded protein amino acids^61,111^. Here, to capture such sequence specificity, we employ a novel reparametrization^126^ of the high-resolution HPS model from Mittal group^103^ which accounts for sequence-dependent hydrophobic and cation–*π* interactions by means of short-range pairwise potentials, and for electrostatic interactions through Yukawa long-range potentials (see section *SI* of the Supplementary Information (SI)). Bonded interactions between subsequent amino acids are restrained by a harmonic potential (Eq. S2), and non-bonded hydrophobic interactions are modelled via an Ashbaugh-Hatch potential (Eq. S4). Additionally, cation–*π* and electrostatic interactions are described by Lennard-Jones (Eq. S5) and Yukawa/Debye-Hückel potential terms (Eq. S3) respectively. The low salt physiological concentration regime (~ 150 mM) of the implicit solvent is controlled by the screening length of the Yukawa/Debye-Hückel potential. Given that the original HPS model^103^ has been shown to underestimate LLPS-stabilising cation–*π* interactions^85^, we employ the recently proposed reparameterization by Das *et al.*^126^. Additionally, to account for the ‘buried’ amino acids contained in the protein globular domains, we scale down those interactions with respect to the original set of HPS parameters by a 30% as proposed in Refs.^85,127^. All the details regarding the model parameters and simulation setups are provided in the Supplementary Information.

To validate the model^103,126^, we evaluate the relative ability to phase separate of several archetypal RNA/DNA-binding proteins that are known to undergo LLPS both *in vivo* and *in vitro*. These proteins are: fused in sarcoma (FUS),^38–40^, heterogeneous nuclear ribonucleoprotein A1 (hnRNPA1)^15,16^ and the TAR DNA-binding protein 43 (TDP-43)^41,42^ (Fig. 1A). We evaluate the phase diagram for either the full protein sequences as well as for some of their specific protein domains such as the RNA recognition motifs (RRMs) or the prion-like domains (PLDs). More precisely, we focus on the following sequences: FUS (full-sequence), FUS-PLD, hnRNPA1 (isoform A1-B, and hereafter named as hnRNPA1), hnRNPA1-PLD, hnRNPA1-RRM, TDP-43 (full-sequence), and TDP-43-PLD (sequences are provided in the SI). For TDP-43, we also distinguish between two different variants, one including the *α*–helix structured domain in the C-tail intrinsically disordered region (h-TDP-43), and another in which the whole PLD region remains fully disordered (wt-TDP-43). Despite h-TDP-43 and wt-TDP-43 only differing by less than 10 % of their sequence structural conformation^128,129^, the presence of the *α* helical structured domain has been shown to moderately affect the protein ability to phase separate^130^. We also study the low complexity domain (LCD) of the isoform A1-A of hnRNPA1^15^ (termed as hnRNPA1-A-LCD), since it has been shown to be a key part of the hnRNPA1 sequence in promoting LLPS in absence of RNA^15^.

**FIG. 1.**
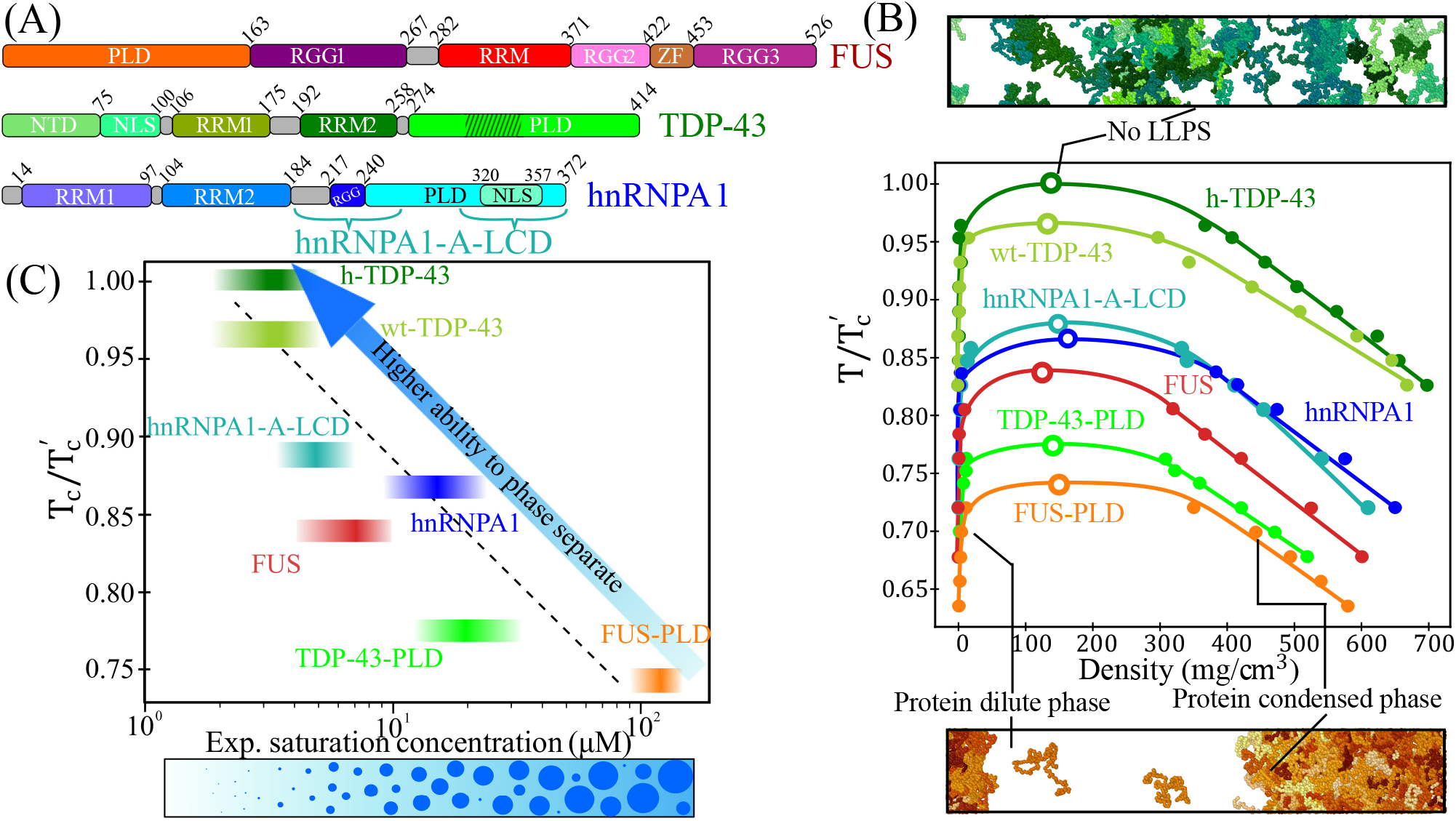
Experimental validation of the sequence-dependent coarse-grained model. (A) Different sequence domains of the three studied proteins, FUS, TDP-43 and hnRNPA1: Prion-like domain (PLD), Arginine-Glycine-rich regions (RGG), RNA-recognition motifs (RRM), zinc finger (ZF), N-tail domain (NTD) and nuclear localization sequence (NLS). Dashed lines in TDP-43 PLD indicate the position of the *α*–helical domain. Braces in hnRNPA1 sequence indicates the two LCD segments composing the sequence of the isoform A1-A of hnRNPA1 (hnRNPA1-A-LCD), corresponding to the residues 186-251 + 304-372 (see section *SII* for the sequences) (B) Phase diagram in the 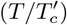-density plane for FUS (red), FUS-PLD (orange), hnRNPA1-A-LCD (turquoise), hnRNPA1 (blue), h-TDP-43 (dark green), wt-TDP-43 (lime green) and TDP-43-PLD (light green). Filled circles indicate the coexistence densities obtained through DC simulations, and empty circles the estimated critical points via the critical exponent and rectilinear diameter laws^134^ (Eqs. (S6) and (S7) in the Supplementary Information). 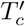 accounts for the highest critical temperature of the protein set (h-TDP-43), which is 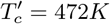. Top: DC simulation of h-TDP-43 above the critical point where LLPS is no longer observed. Bottom: Direct Coexistence simulation of FUS-PLD at 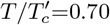 exhibiting two coexisting phases. In both snapshots, different protein replicas are represented by distinct colour tones. (C) Experimental saturation concentration of the proteins to undergo phase separation *versus* the renormalized critical temperatures shown in (B). The experimental saturation concentration^136^ at physiological salt conditions for FUS^50,61,85^ (including FUS-PLD), hnRNPA1^15^, hnRNPA1-A-LCD^137,138^, TDP-43^42,85^ and TDP-43-PLD^135^ are depicted by intervals to consider concentration uncertainty. The height of the intervals accounts for the computational uncertainty in 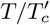. The dashed black line is a linear fit to the displayed data (considering the mean concentration and critical temperature of the interval) and the blue arrow indicates higher ability to phase separate. At the bottom, a schematic cartoon summarizing the expected phase behavior while increasing protein concentration is included. Note that temperatures in this model are unrealistic and only describe the relative ability of the different proteins to phase separate, thus, temperature is only meaningful when is renormalized.

By means of Direct Coexistence simulations (DC)^131–133^ in combination with the laws of rectilinear diameters and critical exponents^134^, we compute the phase diagram (Fig. 1B) of all the aforementioned proteins (hnRNPA1-PLD and hnRNPA1-RRM are shown in Fig. S3 of the SI; see *SIII* section of the SI, or Ref.^101^ for details on how to extract coexisting densities from DC simulations). The predicted renormalized critical points (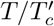 where 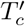 refers to the highest critical temperature of the set) against the experimental saturation concentration of the proteins to undergo LLPS in FUS^61,85^, FUS-PLD^61^, hnRNPA1^15^, hnRNPA1-A-LCD^15^, wt-TDP-43^42,85^, h-TDP-43^42,85^ and TDP-43-PLD^135^ is plotted in Fig. 1C (please note that the experimental saturation concentration reported by Molliex *et al.*^15^ corresponds to the isoform A1-A, but the difference in the critical concentration between the two isoforms is assumed to be minor). We find a positive correlation between the predicted critical point in our simulations and the experimental protein saturation concentration at physiological salt concentration. The uncertainties in the determination of both the critical temperature and the experimental saturation concentration in Fig. 1C are depicted by the height and the width of the coloured bands, respectively. Such impressive qualitative agreement (coarse-grained models with implicit solvent are not expected in principle to quantitatively capture the actual *T_c_*) demonstrates that the Cation-*π* reparameterization proposed by Das *et al.*^126^ on top of Mittal’s group model^103^ is able to describe the relative ability of these proteins to self-assemble into phase-separated condensates with better agreement than the original HPS model^103^ (Fig. S2). Furthermore, we observe a non-negligible difference between the phase diagram of the *α*–helical structured TDP-43 and that of the wt-TDP-43, showing the latter a moderately lower critical temperature, as reported in Ref.^130^. Notably, both prion-like domains of FUS and TDP-43 exhibit a significant lower ability to phase separate than their full counterparts as experimentally found^61^. On the contrary, hnRNPA1-A-LCD exhibits a similar critical temperature than that of the hnRNPA1 full-sequence^15^. To rationalise these observations, in the following section, we perform a detailed molecular and structural characterisation of the condensates.

### B. Structural and interfacial properties of the condensates without RNA

The specific composition and patterning of the amino acids along the sequence has a huge impact on the protein macroscopic phase behavior^86,103,129^. Moreover, beyond sequence, the protein conformational ensemble plays a crucial role not only in their ability to phase separate^116^, but also in the condensate structure^129,130,139,140^. A close example of this is TDP-43, in which a subtle conformational difference on its C-terminal intrinsically disordered domain produces a moderate change on its phase diagram (Figure 1B). To further characterize the molecular, structural and interfacial properties of the previous protein condensates, we now perform a comprehensive full analysis of their surface tension, LLPS-stabilising most-frequent contacts, protein conformational ensembles in and out of the droplet, and condensate structure.

In Fig. 2A we plot the surface tension (*γ*) between the condensate (protein-rich) and protein-poor liquid phases as a function of temperature (renormalized by the highest critical temperature of the protein set, 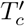 of h-TDP-43). An advantage of computer simulations is that *γ* between two coexisting fluid phases (or between a fluid and a vapour one) can be easily computed, as explained in *SIV*, compared to more challenging approaches (i.e., based on the tie-line width of the phase diagrams) as required in experimental setups^53,141^. We find that the conformational difference in the 40-residue helical region of the TDP-43-PLD terminal domain has significant consequences on the droplet surface tension of TDP-43. For the whole range of studied temperatures, wt-TDP-43 shows smaller *γ* than h-TDP-43. At the same temperature, the presence of the helical structure in h-TDP-43 promotes a more compact assembly of proteins in the condensed phase, increasing the surface tension. Additionally, TDP-43-PLD droplets present much smaller *γ* than those of any of its two full-sequence variants at moderate temperatures, explaining why TDP-43-PLD domains are markedly exposed towards the interface in wt-TDP-43 condensates (Fig. 2B). Similarly, the surface tension of FUS-PLD droplets is lower than that of FUS (full-sequence). However, interestingly, *γ* for hnRNPA1 and hnRNPA1-A-LCD droplets is remarkably similar (as their phase diagrams, see Figs. 1B and S3), confirming the significant importance of the hnRNPA1-A-LCD sequence in contributing to phase separation^15^. Our results clearly evidence a direct correlation between droplet surface tension and condensate stability except for wt-TDP-43 and h-TDP-43 condensates where their characteristic heterogeneous arrangement contributes to decrease *γ* (Figure S5B). Proteins with higher *γ* can typically phase separate until higher temperatures or at lower protein concentration.

**FIG. 2.**
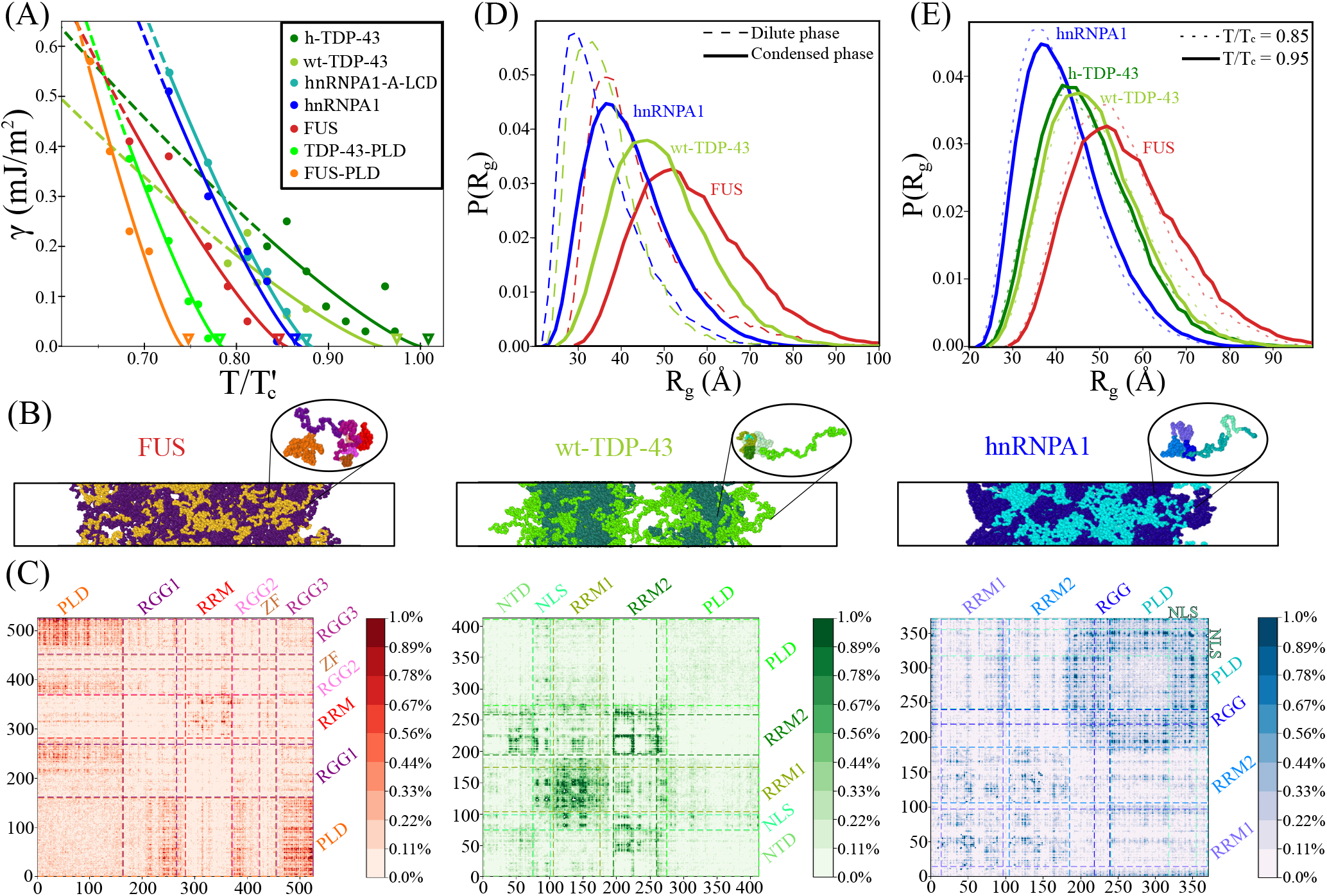
Molecular, structural and interfacial properties of different RNA-binding protein condensates in absence of RNA. (A) Condensate surface tension (*γ*) of FUS, FUS-PLD, hnRNPA1, hnRNPA1-A-LCD, wt-TDP-43, h-TDP-43 and TDP-43-PLD as a function of temperature (renormalized by the highest critical temperature of the protein set, 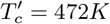 for h-TPD-43). Filled circles indicate the obtained *γ* from DC simulations (see section *SIV* in the Supp. Info. for further details on the calculations) and solid curves the *γ* ∝ (*T_c_ - T*)^1.26^ fit to our data^134^ (dashed curves depict the predicted surface tension at low T extrapolated from the fit). Empty down triangles represent the obtained (renormalized) critical temperatures of each sequence using the law of rectilinear diameters and critical exponents as in Fig. 1B. (B) Snapshots of Direct Coexistence simulations of the three full sequences at *T* = 0.9*T_c_* (meaning *T_c_* the critical temperature of each protein): FUS (left, *T* = 360*K*), wt-TDP-43 (center, *T* = 410*K*) and hnRNPA1 (right, *T* = 390*K*). FUS, wt-TDP-43 and hnRNPA1 prion-like domains are highlighted in orange, bright green and cyan respectively, while the rest of their sequences in purple, dark green and dark blue respectively. The structure of the condensates clearly show the contrast between homogeneously distributed PLD domains as in FUS, clustered PLD domains as in hnRNPA1, or interfacially exposed PLD domains as in wt-TDP-43 condensates. (C) Frequency amino acid contact maps of FUS (left), wt-TDP-43 (center) and hnRNPA1 (right) droplets at *T* = 0.9*T_c_*. Scale bars indicate the averaged percentage of amino acid contact pairs per protein (see section *SV I* of the Supp. Info. for further details on these calculations). Dashed lines depict the limits of the different protein domains as indicated in Fig. 1A. (D) Protein radius of gyration distribution function of the three sequences at *T/T_c_* = 0.9 and at the bulk equilibrium coexisting density of the diluted (dashed curves) and the condensed phase (continuous curves). (E) Protein radius of gyration distribution function within the condensates at moderate (*T/T_c_* = 0.85) and high temperature (*T/T_c_* = 0.95).

Next, we focus on the structural organization of the different protein condensates. A significant contrasting behavior between both FUS and hnRNPA1 droplets and those of TDP-43 (both variants) is observed. While both FUS and hnRNPA1 exhibit homogeneous density droplet distribution with their PLDs indistinctly located along the condensate (although in hnRNPA1 more clustered), TDP-43 shows a highly marked heterogeneous distribution exposing its prion-like domains towards the droplet boundaries (Fig. 2B), evidencing that their PLD interactions barely favor aggregation^129,130^. This condensate arrangement allows the minimisation of the droplet surface tension and the simultaneous maximisation of its enthalpic gain (in absolute value) through a higher density of LLPS-stabilising contacts at the droplet core^142^. In the case of wt-TDP-43, such structural heterogeneity is so pronounced, that condensates split into smaller nearly-interacting liquid droplets as shown in Fig. 2B (center). Conversely, the *α*-helix structure of h-TDP-43 notably favors the interaction between helical domains, and hence, between the rest of the intrinsically disordered neighbour regions significantly enhancing the PLD connectivity, and thus, reducing droplet heterogeneity as experimentally suggested^129^. Moreover, our simulations show that the structured *α*–helical domain considerably reduces the local density fluctuations of the droplet and further stabilises the condensate (Fig. 1B).

To rationalise the molecular driving forces behind these structural differences, we compute: 1) the amino acid contact map frequency of the proteins within the condensates (Fig. 2C and S8); and 2) the most persistent residue-residue pair interactions along the aggregated proteins (Fig. S9-11). We develop a smart cut-off analysis of each specific residue-residue interaction (adapted to the range of the HPS potential^103^, see section *SV I* of the Supp. Info. for further details) to elucidate the key molecular interactions promoting LLPS according to our model^103,126^.

In FUS condensates, the most repeated contacts are G-G, R-Y and G-Y (Fig. S9A), highlighting how hydrophobic, cation–*π*, and more modestly electrostatic interactions contribute to stabilise the droplets. Since Glycine (G) represents nearly the 30% of the residues along FUS sequence, the frequency of G-G is the highest despite not being one of the strongest pair of amino acid interactions^85^. However, when normalizing the computed number of contacts by the amino acid abundance, we find that the cation–*π* interaction R-Y becomes the most relevant one inducing LLPS^76,77^ according to this force field (see Fig. S9B). Furthermore, when analysing the FUS contact map (Fig. 2C), we observe that its prion-like domain, despite showing a much lower ability to phase separate on its own than the full protein, markedly interacts with the three RGG domains. The top contacts of the PLD alone are very different from those of the full-sequence FUS (Fig. S9A), resulting in much worse phase separation capabilities for the PLD than for the full-FUS sequence (Fig. 1B) as experimentally observed^50,61^. We also find moderate LLPS-stabilising interactions among different RNA-recognition motifs in FUS (Fig. 2C).

While in FUS condensates the PLD plays a vital role in LLPS^39,103^, the aggregation of TDP-43 (wild-type) is mainly sustained by contacts between RRMs, either with themselves or with other protein regions such as the NTD or the NLS, but mostly dominated by RRM1-RRM1 interactions (Fig. 2C). Nonetheless, the wt-TDP-43 PLD region is still the second protein domain establishing more contacts in total after the RRM1 segment, but mostly because of its length. The three most predominant contacts in wt-TDP-43 (according to our model^103,126^) are K-F, K-E and K-D (Fig. S10A), clearly denoting the key role of cation–*π* and electrostatic interactions in driving condensation. However, when the structured helical region is present (h-TDP-43), R-F contacts sensibly increase becoming the third most dominant interaction. Interestingly, the renormalization of contacts by amino acid abundance in TDP-43 barely modifies the list of the most frequent interactions, probably due to the very homogeneous distribution of amino acids along its sequence (Fig. S10C) when compared to that of FUS. However, similarly to FUS, TDP-43-PLD shows a completely different list of the most repeated interactions compared to the full protein (Fig. S8A), which is likely contributing to reduce its critical temperature (Fig. 1B) and surface tension (Fig. 2A).

In hnRNPA1, the most frequent contacts are G-G, G-S, and G-R (Fig. S11A), but since glycine is the most abundant amino acid (~ 25%) followed by serine (~ 15%), the normalized contacts by amino acid abundance show that R-Y, R-F and K-Y are dominant interactions, again highlighting the importance of cation–*π* interactions in hnRNPA1 LLPS. The list of top interactions of hnRNPA1-PLD, even after normalization, is very similar to that of hnRNPA1 (Fig. S11A-B), which explains why the phase diagrams of both sequences are hardly distinguishable (Fig. S3A). Surprisingly, the list of the most frequent interactions of hnRNPA1-A-LCD is also remarkably similar to that of hnRNPA1 full-sequence (Fig. S11A). In fact, the detailed contact map of hnRNPA1-A-LCD corresponds to the region of hnRNPA1 that contains more LLPS-stabilising interactions (dashed lines in Fig. S8). Thus, the ability of hnRNPA1 to phase separate alone can be mainly captured by the present protein interactions in hnRNPA1-A-LCD (see Fig. S3A).

Finally, we investigate the protein conformational ensemble within the condensates and the diluted phase by computing the radius of gyration distribution function of the proteins *P* (*R_g_*). Our simulations reveal that in all cases, when proteins transition from the diluted to the condensed phase, their conformations adopt larger radii of gyration (Figure 2D). Also, the width of *P* (*R_g_*) considerably increases, indicating the more versatile conformations that proteins can exhibit within the condensate. This structural behavior allows proteins to maximize their number of intermolecular contacts, and thus, the droplet connectivity as recently shown in Ref.^116^. Phase-separation driven expansion of proteins undergoing homotypic LLPS has been observed for tau-IDP^143^ using steady-state fluorescence measurements of pyrene and fluorescein-labeled tau-K18 proteins, a protein associated with Alzheimer disease^62^. Even if modest, phase-separation induced expansion enables IDRs to establish a surplus of enthalpy-maximizing (more energetically favourable) inter-protein contacts in the condensed phase, compared to those that they would adopt if they remained unchanged or underwent collapse. On the other hand, very recently, NMR and EPR spectroscopies have shown that the N-Terminal domain of FUS is compacted when entering in the condensed phase under agarose hydrogel conditions^144^. However, due to the employed different experimental matrix composition, our model predictions cannot be directly related to these striking observations. Now, when regarding to the protein conformational ensemble within the condensates along temperature, we note a mild change in the hnRNPA1, FUS and TDP-43 conformations as we approach to the critical *T* (Figure 2E), in contrast to those measured in the diluted phase as a function of *T* (Fig. S6 and Ref.^86^). Moreover, when comparing both TDP-43 variants *P* (*R_g_*) distributions, we find almost identical protein ensembles, exhibiting the wild-type variant slightly more open conformations. Such small surplus of extended conformations shown by wt-TDP-43, which can enable a higher amount of intermolecular contacts^116^, is not enough to enhance LLPS as through the *α — α* helical interactions present in the h-TDP-43^129^.

### C. RNA-induced reentrant behavior in phase separation

RNA has been recently shown to critically regulate both the phase behavior of different RNA-binding proteins^43,50,52,56,57,145^, and most importantly, the emergence of aberrant liquid-to-solid pathological phase transitions^51,62^. In this section, we explore the impact of poly-U RNA in LLPS of RBPs from a molecular and a physico-chemical perspective. By means of the novel coarse-grained model of RNA recently proposed by Regy *et al.*^119^ and Direct Coexistence simulations^131–133^, we characterize the condensate stability of different RNA-binding proteins (and domains) from low to moderately high poly-U concentration regimes. We choose poly-U RNA for simplicity^85^ and to follow previous landmark works on RNA-RBP phase separation^43,56^.

First, we mix poly-U RNA strands of 250 nucleotides with the proteins studied above. Remarkably, not all proteins were able to favorably interact with poly-U in our simulations. We find that FUS-PLD and TDP-43 (including both variants) do not associate with poly-U even at very low RNA concentration (i.e., ~ 0.05 mg poly-U/mg protein). We further test the affinity of wt-TDP-43 with poly-U strands by performing a separate analysis of each of its major protein sequence domains (PLD, RRM1 and RRM2). None of these domains exhibited a conclusive interaction with poly-U at temperatures moderately below the critical one. That is not entirely surprising since: 1) Several experimental studies have shown that TDP-43-RRM1 only presents a strong affinity for RNA strands enriched in UG nucleotides^146–148^, and 2) TDP-43-RNA heterotypic interactions are mainly driven by the RRM1, whereas the RRM2 plays a supporting role^146^. Furthermore, in the employed model, the interactions between poly-U and TDP-43 are mainly electrostatic, and therefore, other factors such as RNA secondary and tertiary structures that might sensibly promote RRM binding to specific RNA sequences are not explicitly considered^149^. On the contrary, the non-interacting behavior between FUS-PLD and poly-U strands was completely expected since the FUS-PLD sequence does not present neither RNA-binding domains nor positively charged domains, thus, precluding their association.

We now evaluate the phase diagram of all proteins (or protein domains) that favorably interact with poly-U, these are: FUS, hnRNPA1, hnRNPA1-PLD, hnRNPA1-A-LCD and hnRNPA1-RRMs. In all these systems, except for hnRNPA1-PLD, the resulting phase behavior is similar to that shown in Fig. 3A-B for FUS (note that poly-U/hnRNPA1-PLD condensates show a very mild LLPS enhancement at low poly-U concentration, Fig. S4 and Table S3, so hereafter, the results are just discussed for hnRNPA1-A-LCD, FUS, hnRNPA1 and hnRNPA1-RRMs). At low poly-U/protein ratios, the stability of the condensates moderately increases (~ 2% higher critical temperature), while at high concentration, the critical point decreases below the critical temperature without RNA (Fig. 3E). This reentrant behavior has been experimentally observed for synthetic peptides such as *RP*_3_ and *SR*_8_ in poly-U mixtures^56^ and for RNA-binding proteins such as FUS^50,55–57^, Whi3^58^, G3BP1^44^, or LAF-1^43^. In fact, for FUS and hnRNPA1 proteins, it has been reported that at RNA/protein mass ratios close to 0.9, phase separation can be inhibited^57^, which is in qualitative agreement with our observations (~ 0.3 mg RNA/ mg protein). The higher RNA reentrant concentration measured *in vitro* may come from the fact that it refers to the total solution concentration rather than within the phase-separated condensates, as in our simulations, which is very likely to be lower than in the diluted phase. From our simulations, we also note that although a 2% shift in the critical temperature might seem insignificant, the actual increment in temperature according to the force field^103,119,126^ may be as large as 10K, which represents a huge temperature rise when referred to the physiological cell environment. We also plot the phase diagram for FUS with poly-U in the RNA/protein mass ratio-density plane for different temperatures close to the pure FUS critical one (Fig. 3C). At the pure FUS critical temperature, we observe a closed-loop diagram (green curve), and for slightly lower temperatures, reentrant phase behavior is also recovered in agreement with experimental findings^50,55–57^. To microscopically rationalize this behaviour, we compute the protein-protein, protein-RNA, RNA-RNA and total number of contacts as a function of poly-U concentration (Fig. S14), which clearly shows that at low RNA concentration, the total number of contacts per protein within the condensates is higher (~ 30) than at high poly-U concentration (~ 20) (just before phase-separation is no longer possible). Moreover, a maximum in FUS-poly-U contacts can be seen at moderate concentration (~ 0.17 mg poly-U/ mg FUS), whereas RNA-RNA contacts are almost negligible at any concentration. We note that to accurately determine the specific RNA-induced temperature raise, atomistic simulations would be needed^150^, although that is far beyond the current computational capability. Nevertheless, just the fact that a coarse-grained model successfully captures the experimental reentrant behavior observed in some RBP-RNA condensates is outstanding^119^. For the studied proteins, FUS (red) exhibits the highest variation in critical temperature at either low and high RNA concentration (Fig. 3D). Interestingly, hnRNPA1 (blue) shows an intermediate behavior between that of its A-LCD (cyan) and RRM (purple) domains. The maximum critical temperature in hnRNPA1-RRM is reached at the lowest RNA concentration of the set and it sharply decays after the maximum. Contrarily, hnRNPA1-A-LCD suffers only a moderate increment of the critical temperature, but its reentrant behavior is smoother and appears at much greater concentration (2 times higher) than that of hnRNPA1-RRM. Overall, hnRNPA1 condensates present higher RNA-induced stabilization in the low RNA regime than that of its PLD and RRMs. Moreover, it is worth mentioning that in all sequences, the larger enhancement of LLPS is reached at a poly-U concentration close to the electroneutrality point (depicted by crosses in Fig. 3D), which emphasizes the major importance of electrostatic nucleotide-amino acid interactions in RNA-RBPs phase separation^56,119^.

**FIG. 3.**
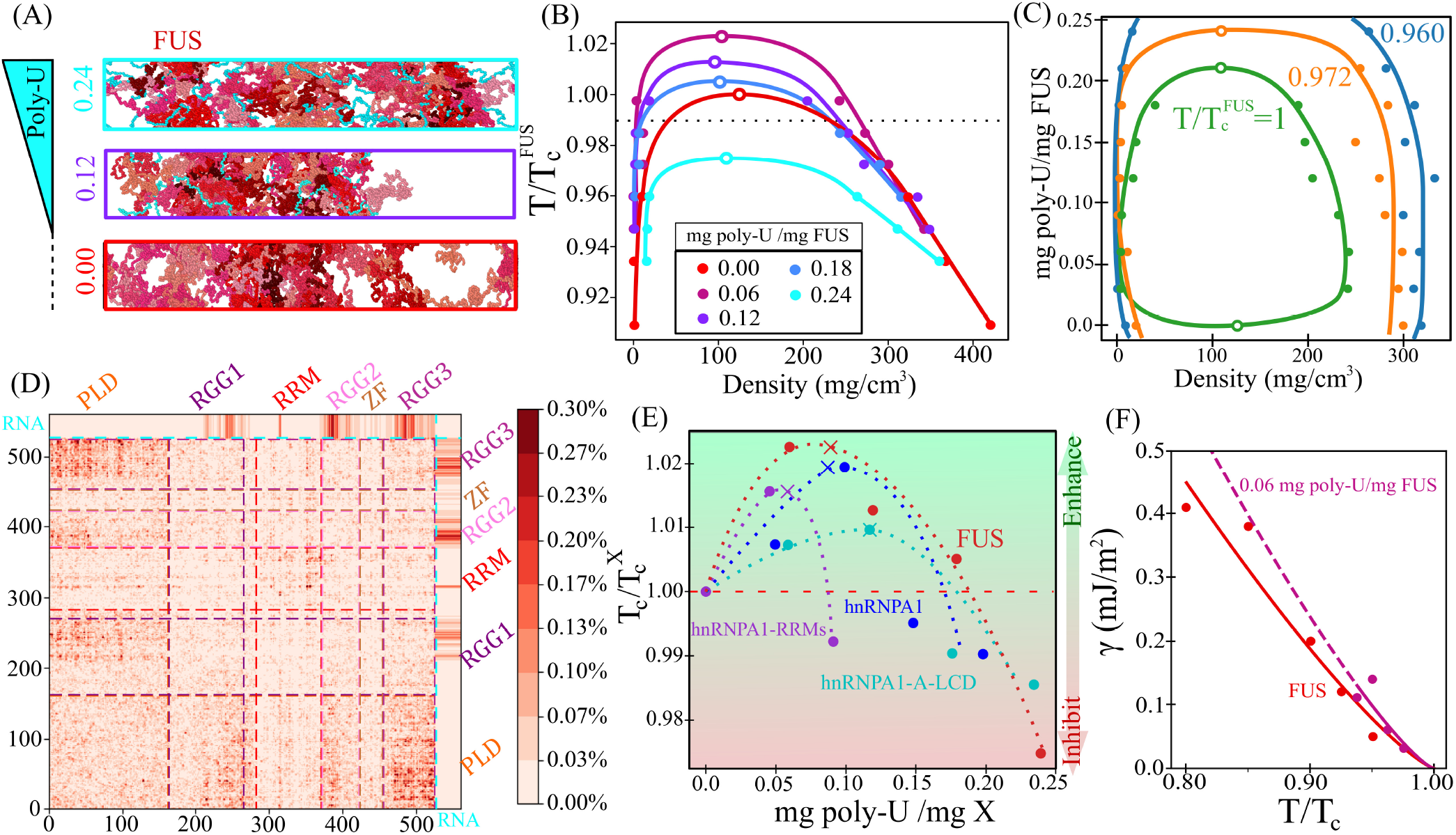
RNA-induced reentrant behavior in RBP phase separation. (A) Snapshots of Direct Coexistence simulations of FUS (red) and poly-U (cyan) at temperature (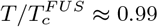, where 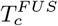 refers to the critical temperature of FUS in absence of poly-U) with increasing poly-U/FUS mass ratios as indicated at the left side of the simulation boxes. (B) Phase diagrams in the temperature-density plane for five different poly-U/FUS mass ratios as indicated in the legend. Empty circles represent the estimated critical point and filled circles the obtained coexisting densities from DC simulations. The horizontal dotted line depicts the temperature at which the DC snapshots shown in (A) were taken. (C) Phase diagram in the (poly-U/FUS mass ratio)-density plane for three different temperatures: 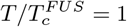 green, 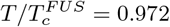 orange, and 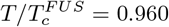 blue. Empty circles depict the estimated critical points while filled circles the obtained coexisting densities from DC simulations. (D) Map of intermolecular contacts per protein replica for FUS in presence of poly-U(250) at *T/T_c_* = 0.9 and at the coexisting droplet equilibrium density at that temperature. The ratio between poly-U(250nt) and FUS is 0.119. The intermolecular contacts between poly-U RNA and FUS are included in the upper and right side edges of the map. Distinct domains of FUS have been labeled as in Fig. 2 C (Left). (E) Reentrant behavior of several RNA-binding proteins/domains as a function of the poly-U/protein mass fraction. Filled circles depict the critical temperature (renormalized by that of each pure protein in absence of poly-U) of the different protein mixtures. Cross symbols indicate the poly-U/protein mass fraction at which mixtures possess neutral electrostatic charge, and the horizontal red dashed line shows the limit above which phase separation is enhanced by poly-U (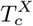 is the critical temperature of each pure protein/domain). (F) FUS droplet surface tension (*γ*) as a function of temperature (renormalized by *T_c_* of each system) with (purple) and without (red) poly-U as indicated in the legend. Filled circles account for the obtained *γ* values from DC simulations, whereas solid lines account for the fit given by the following expression^134^ *γ* ∝ (*T − T_c_*)^1.26^, which can be conveniently extrapolated to moderate lower temperatures (dashed curve).

To characterize the RNA-RBP condensates from a microscopic perspective, we analyze the key molecular contacts enabling phase separation (Figs. 3 D, S12 and S13). We find that, near the optimum poly-U/protein concentration promoting LLPS, the most frequent contacts in poly-U/FUS condensates are now R-U and G-U (Fig. S13). This demonstrates how poly-U (at low fraction) plays a major role in sustaining the condensates, given that the two most frequent contacts are now shifted from G-G and R-Y, to the electrostatic cation-anion R-U interaction; and G-U interactions (Fig. S13). In terms of the FUS sequence domains, the RGG regions and the RNA-recognition motif are those presenting more contacts with the poly-U strands, explaining why G-U becomes one of the most dominant molecular contacts by proximity (Fig. S12). On the other hand, the PLD region presents the least favorable interactions with poly-U. The fact that poly-U strands are not specifically recognized by the ZF domain needs to be further tested in order to check whether that may be caused by model deficiencies (lack of secondary- or tertiary-driven interactions), and/or due to the fact that poly-U strands are not specifically recognized by ZF domains^122,151^.

We also analyze the protein and RNA conformational ensemble as a function of poly-U concentration by computing the radius of gyration histograms for FUS and poly-U (125nt) (Figure S7). We strikingly find that despite varying the stability and density of the droplets with RNA concentration (Figs. 3A, B, C and E), the structural conformation of the proteins and RNA does not significantly change (at least by analyzing the *R_g_*). Regarding poly-U/hnRNPA1 droplets, our simulations reveal that G-G contacts remain as the dominant amino acid pair interaction (although it substantially decreases by a factor of two), and R-U and G-U become the next two following most frequent contacts (further details in section *SV I* and Fig. S13). However, the behavior of poly-U/hnRNPA1-A-LCD condensates is radically different, despite its phase diagram being altered by poly-U addition the most frequent contacts remain similar to those in absence of RNA but including a very modest excess contribution in R-U interactions (Fig. S13). On the contrary, when just considering the RRM1-RRM2 hnRNPA1 domains (purple curve in Fig. 3E), even at the lowest RNA-protein ratio where the droplet stability attains its maximum value, R-U and K-U emerge as some of the most frequent contacts in spite of the very modest poly-U concentration (Fig. S13). Finally, if we examine the contact map between poly-U and different hnRNPA1 (full-sequence) domains, we strikingly observe that the PLD comprises the highest amount of interactions with poly-U strands. However, such observation can be explained through the longer length of the PLD with respect to the two RNA-recognition motifs. Yet, the strongest electrostatic interactions (mainly R-U and K-U) between hnRNPA1 and poly-U are those held through the two RRM domains (Fig. S12).

We also determine the surface tension (*γ*) of the condensates with the dilute phase in presence of poly-U as a function of temperature (Fig. 3F). Both for FUS (Fig. 3F) and hnRNPA1-A-LCD (Fig. S5) condensates, we observe that poly-U at low concentration significantly increases the droplet surface tension besides further stabilizing the droplets as shown in Fig. 3D. Our simulations suggest that the molecular origin behind such surface tension increase comes from the reallocation of the positively charged residues (R, H, and K) within the bulk condensate to maximize the molecular connectivity with poly-U, rather than remaining more exposed to the interface as in the pure component, and therefore, contributing to minimise the droplet surface tension due to their higher hydrophilicity. On the contrary, at moderately high poly-U ratios, the surface tension seems to decrease, although the scattering of our data does not allow us to conclude whether a non-monotonic behavior in *γ* may also exist (Fig. S5).

To further elucidate the role of RNA-regulated RBPs condensate stability, we now focus on the effect of poly-U length in LLPS. A landmark study by Maharana *et al.*^57^ showed that smaller RNAs were more potent than larger ones in solubilizing FUS condensates. On the other hand, Zacco *et al.*^60^ found that longer RNA repeats presented weaker dissociation constants with N-RRM1-2 domains of TDP-43 than 3-fold shorter RNA strands. Given the critical role that RNA performs on the behavior of many different RBP organelles^15,43,58^, we investigate the role of RNA length by introducing poly-U strands of different lengths (i.e., 10, 50, 100, 125 and 250 nucleotides) at a constant poly-U/protein mass ratio that maximises droplet stability (~ 0.12 mg RNA/mg protein) for FUS and hnRNPA1-A-LCD sequences (Fig. 3E). Our simulations reveal that very short poly-U strands (~ 10 nt) do not enhance phase separation in FUS and hnRNPA1-A-LCD droplets (Fig. 4A-B). In fact, 10 nt poly-U strands in hnRNPA1-A-LCD droplets inhibit LLPS even at low concentration. On the other hand, we observe that RNA strands longer than ~100 uridines (hereafter called minimum critical length) promote a similar droplet stabilization independently of their length (Fig. 4B). To further investigate the molecular insights behind these observations, we analyze the FUS/RNA conformational ensemble within phase-separated droplets with distinct RNA lengths by computing their radius of gyration histograms. As it can be seen in Fig. 4C, RNA strands with radii of gyration comparable or longer than those of the proteins (i.e., above the minimal critical length, Fig. 4B) promote maximum condensate stabilization, whereas RNA poly-U strands with shorter *R_g_* than those of FUS proteins (i.e., below the critical length) cannot achieve the same degree of droplet stabilization (and density) for the same FUS/RNA concentration. We note that the observed minimum critical RNA length in FUS and hnRNPA1-A-LCD droplets may be also modulated by some protein/RNA specific features/modifications such as RNA sequence, secondary structure interactions, protein charge distribution, post-translational modifications or RRM patterning effects^43,51^. Moreover, if we compute the number of protein contacts within the condensate when adding short and long RNA chains (Fig. S14B), we find that strands longer than the minimum critical length promote higher amount of protein intermolecular interactions, while short RNA chains (i.e., 10 nt) considerably hinder the liquid-network connectivity of the proteins within the droplets^117^, hence, RNA behaving as a ligand/client instead of a co-scaffold, as when RNA is longer than 100 nucleotides. An extensive characterisation (and rationalization) of the critical aspects controlling RBP-RNA aggregation, such as the RNA length dependence studied here, may provide highly valuable insights for designing therapeutic RNA strategies to combat neurodegenerative disorders whose development is deeply linked to aberrant accumulation and solidification of RBP condensates^63,138^.

**FIG. 4.**
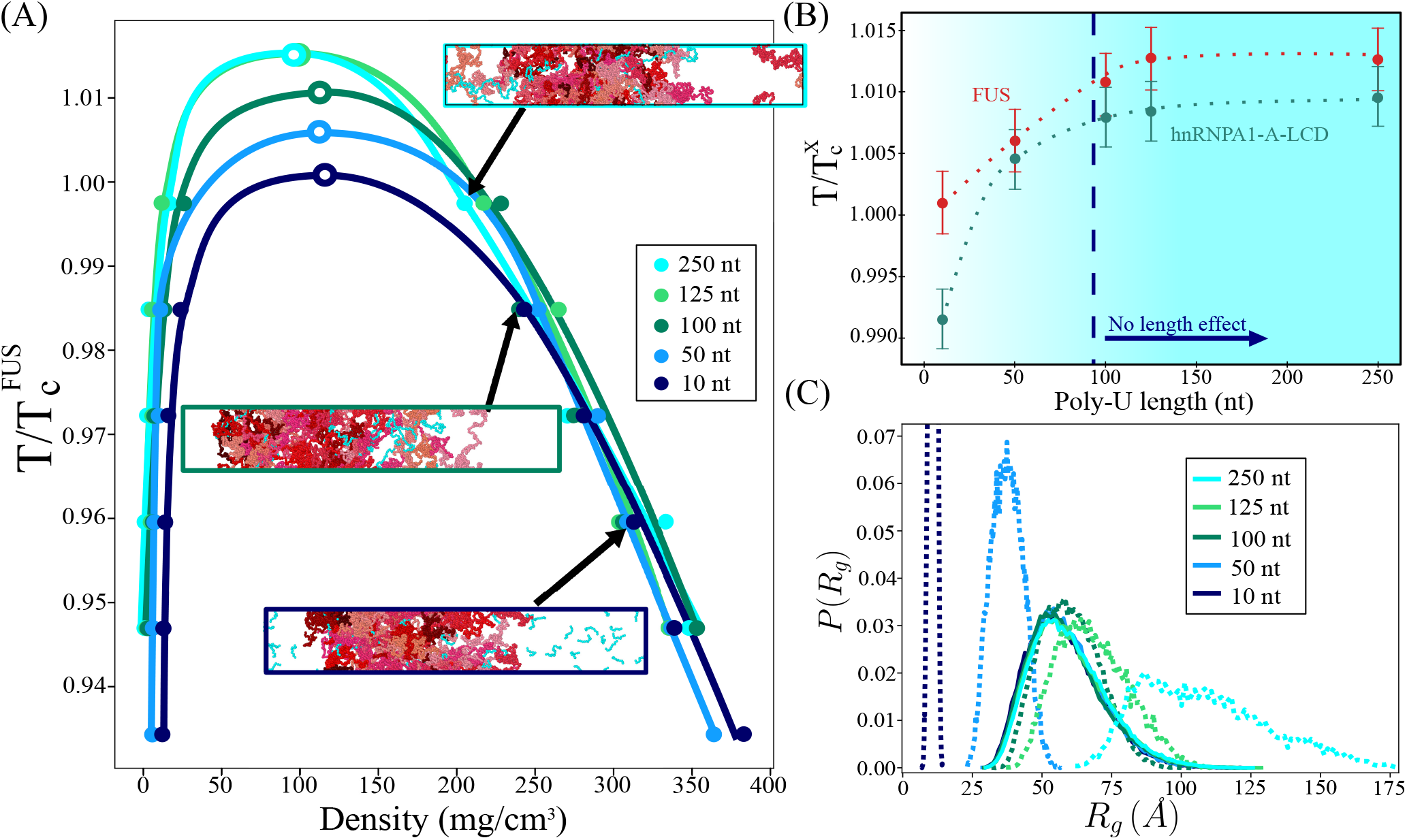
Condensate stability dependence on poly-U RNA length. (A) Phase diagrams in the temperature-density plane for poly-U/FUS mixtures of different poly-U strand lengths (as indicated in the legend) at a constant concentration of 0.119 mg poly-U/mg FUS. Temperature is normalized by the critical one of FUS 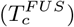 without poly-U RNA. DC snapshots of three representative cases of poly-U/FUS mixtures at the temperature indicated by the arrow and with poly-U lengths as depicted by the box side color (see legend) are also included. (B) Renormalized critical temperature of poly-U/FUS (red) and poly-U/hnRNPA1-A-LCD (green) condensates as a function of poly-U length for a constant concentration of 0.119 mg poly-U/mg FUS and 0.117 mg poly-U/mg hnRNPA1-A-LCD respectively. Temperature is normalized by the corresponding critical temperature 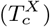 of each protein in absence of poly-U. The vertical dashed line indicates the minimum RNA length required to maximise droplet stability at this given concentration. (C) Radius of gyration histograms of FUS (continuous curves) and poly-U (dotted curves) extracted from the condensed phase of the NVT simulations shown in (A) for different strand lengths as indicated in the legend. Please note that all histograms have been normalized.

### D. RNA modulates the transport properties of RBP condensates

Besides controlling condensate stability, RNA has been proved to play a critical role in regulating the dynamics of many membraneless organelles^15,51,57^. A seminal study of Zhang *et al.*^58^ showed that the RNA-binding protein Whi3 phase separates into liquid-like droplets whose biophysical properties can be subtly tuned by changing the concentration of the mRNA binding partner, showing that larger RNA content increases Whi3 droplet viscosity. On the other hand, RNA has been also observed to provoke the opposite effect in LAF-1 condensates when short strands (50 nt) were introduced^43^. Nonetheless, when long RNAs were used (up to 3,000 nt), LAF-1 condensates presented significantly higher viscosity^49^. Moreover, beyond length, RNA sequence can be also an important factor in modulating droplet dynamics^152^. However, a full understanding of the precise effect of RNA in different RBP condensates still requires further work^54^. Here, we aim to provide new molecular insights on this discussion by measuring via computer simulations the protein diffusion and viscosity of several RBP condensates as a function of poly-U concentration and for different poly-U lengths.

*In vitro*, viscosity (*η*) is usually obtained by bead-tracking within droplets using microrheology techniques^29,43,153,154^, so that the trajectory can be registered and the mean squared displacement (MSD) of the beads calculated, and thus, their diffusion coefficient. Then, the droplet viscosity is inferred from the diffusion coefficient by using the Stokes-Einstein relation^155^. However, in computer simulations we can measure both observables independently. The linear viscoelasticity of a material can be straightforwardly computed by integrating in time the relaxation modulus *G*(*t*) of the system^156,157^ (see Section *SV II*), whereas the diffusion coefficient can be extracted from the MSD of the proteins. The direct calculation of *G*(*t*) provides useful information about the underlying relaxation mechanisms of the proteins (see Fig. 5A for FUS condensates with and without poly-U), either at short times (white region) where the relaxation modes mostly depend on short-range and intramolecular interactions (i.e., internal protein conformational fluctuations such as bond or angle relaxation modes), or at long time-scales (beige region) where *G*(*t*) is dominated by intermolecular forces, long-range conformational relaxation, and protein diffusion within crowded liquid-like environments. Moreover, in Figure 5A, the fact that *G*(*t*) presents a faster decay when condensates contain RNA (purple circles) suggests that their viscosity will be lower than those of pure FUS droplets (red circles).

**FIG. 5.**
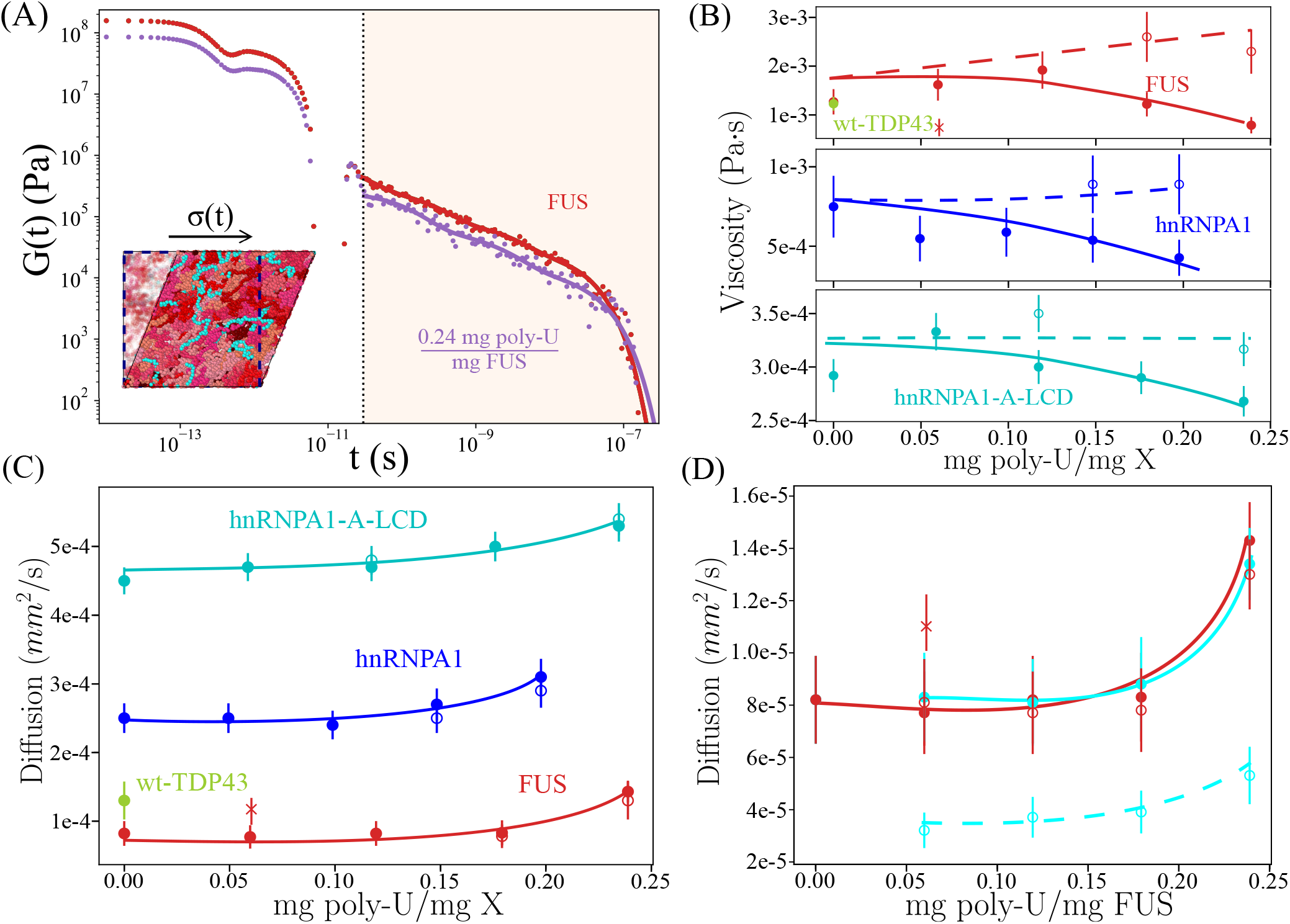
RNA critically regulates the dynamical properties of RBP condensates. (A) Shear stress relaxation modulus of FUS condensates in absence (red) *versus* presence (purple) of poly-U strands of 125 nucleotides at 0.24 mg poly-U/mg FUS mass fraction, *T/Tc* = 0.97 (where *Tc* refers to the critical temperature of FUS pure condensates) and the corresponding equilibrium bulk density of each droplet at such conditions. The vertical dotted line separates the fast-decay relaxation mode regime (white) and the slow-decay relaxation mode one (beige). A snapshot illustrating a shear stress relaxation experiment over a poly-U/FUS condensate simulation box is included. (B) Viscosity of FUS (at *T/Tc* = 0.97), hnRNPA1 (at *T/Tc* = 0.985) and hnRNPA1-A-LCD (at *T/Tc* = 0.98) condensates as a function of the poly-U/protein mass ratio. An estimate of wt-TDP-43 viscosity in absence of poly-U at *T/Tc* = 0.97 is also included (green circle). Filled circles depict viscosities when poly-U strands of 125 nt were used whereas empty circles when strands 250 nucleotides were added. The cross symbol in the FUS panel indicates the viscosity of FUS-poly-U condensates when strands of 10 nt were included. Continuous and dashed lines are plotted as a guide for the eye for strands of 125 and 250 nt respectively. Note that *Tc* refers to the pure component critical temperature of each protein. (C) Protein diffusion coefficient (filled circles) as a function of poly-U(125 nt)/protein(X) mass ratio. Empty circles account for the protein diffusion coefficient when poly-U strands of 250 nt were added, while filled ones for 125 nucleotide RNA strands. The same system conditions described in (B) are applied on these calculations. Continuous curves are included as a guide to the eye. (D) Diffusion coefficient of FUS (red) and poly-U strands (cyan) as a function of the poly-U/FUS mass ratio. Empty circles show the diffusion coefficient of FUS (red) and poly-U strands (cyan) when RNA strands of 250 nt were added, whereas filled circles correspond to values with poly-U strands of 125 nt. The red cross symbols indicates the diffusion of FUS proteins in condensates with poly-U chains of 10 nucleotides. The diffusion of poly-U chains of 10 nt is *D* ~ 3 × 10^*−*3^*mm*^2^/*s*, so that it has been omitted from the panel.

We characterize the condensate dynamics of FUS, hnRNPA1, and hnRNPA1-A-LCD as a function of poly-U concentration at constant temperature (just below the critical *T* of each protein in absence of poly-U, *T/T_c_* ~ 0.98) and at the corresponding bulk droplet equilibrium density corresponding to each poly-U concentration at that temperature. First, we introduce poly-U strands of 125 nucleotides. As shown in Fig. 4B, the phase diagram for a given concentration is not expected to change either by using strands of 125 or 250 nucleotides. For both FUS and hnRNPA1-A-LCD droplets, we observe a mild non-monotonic behavior with a maximum in viscosity at low poly-U ratios (filled circles in Fig. 5B), which might be directly related to the maximum in droplet stability shown in Fig. 3D, or due to a coincidental scattering of our measurements. Nevertheless, at moderate poly-U mass ratios (i.e., > 0.20 mg poly-U/ mg protein), the viscosity of the condensates (using 125 nt RNA strands) is about 30% lower than that without poly-U. On the other hand, a monotonic decreasing trend in viscosity was detected for hnRNPA1 condensates, where almost a ~ 50% drop in *η* is found at high poly-U mass fractions (0.24 mg poly-U/ mg hnRNPA1). Even though the observed maximum in viscosity could be easily related to the reentrant behavior depicted in Fig. 3E, further work needs to be devoted to clarify whether this is a real feature of the model, and ultimately of these RBP-RNA condensates. Furthermore, we investigate how poly-U strands of 250 nucleotides can regulate droplet viscosity at the same concentrations. While poly-U 125 nt strands significantly reduce viscosity at high mass ratios, poly-U 250 nt strands barely varies the condensate viscosity at the same concentrations, except for FUS, where a moderate viscosity increase was detected (empty symbols in Fig. 5B). These observations are in full agreement with those reported for LAF-1 condensates in presence of short^43^ and long^49^ RNA strands. Long RNA chains, even at low to moderate concentrations, can increase droplet viscosity due to their own slow relaxation times. In fact, when very short RNA strands of 10 nt are added in FUS condensates (cross red symbol), the viscosity of the phase-separated droplets is almost two times lower than that of condensates containing 250 nt at the same poly-U/FUS ratio (~ 0.06 mg poly-U/ mg FUS). In Section *SV II* and Tables S4 and S5 we provide the values of *η* for the different RBP condensates as a function of poly-U concentration and length, as well as details on the statistical analysis for estimating the uncertainty of these calculations. We note that our *G*(*t*) values for FUS do not quantitatively match with those of Ref.^74^. That is somewhat expected since our coarse-grained model has been parametrized to describe the radius of gyration^103^ and most frequent molecular contacts^126^ between proteins rather than dynamic properties such as transport properties within the condensates. Nonetheless, the observed behavior with RNA and between the different RBP condensates is expected to qualitatively hold despite the different model approximations (i.e., implicit solvent and amino acids/nucleotides represented by spherical beads).

Finally, we measure the protein diffusion coefficient (*D*) within the condensates for all previous poly-U concentrations and strand lengths of 125 and 250 nt. In all cases, we find a coherent correlation between viscosity and protein mobility, being the latter considerably higher at moderate poly-U/protein ratios than for pure protein condensates (Fig. 5C). Strikingly, protein diffusion hardly depends on poly-U strand length (empty symbols) as viscosity does (Fig. 5B). Only when extremely short RNA chains of 10 nt are added, as those tested in FUS condensates (Fig. 5D cross symbol), protein diffusion noticeably increases. While the shear stress relaxation modulus of the condensates crucially depends on the RNA strand length (longer RNAs imply longer *G*(*t*) relaxation decay), the protein diffusion coefficient does not. The later mainly depends on droplet density (and temperature) and, as shown in Fig. 4 A, condensates densities remain similar either when using strands of 125 and 250 nucleotides. However, when 10 nt strands are added at the same poly-U concentration, the droplet density critically decreases from ~0.28 *g/cm*^3^ (for 125 and 250 nt chains) to ~0.20 *g/cm*^3^. Therefore, our simulations suggest that the condensate dynamics dependence on RNA concentration is intimately related to the droplet density decrease as a function of poly-U concentration and length shown in Figs. 3B and 4A. To better comprehend the underlying mechanism behind such behavior, we also measure *D* for poly-U strands of different lengths (i.e., 125 and 250 nt) within the condensates. In Figure 5D, we observe the severe impact of RNA chain length on its own mobility, as expected. Whereas FUS *D* (and also condensate stability and density) barely depends on the poly-U length (at least between 125 and 250 nucleotides), a two-fold decrease in the RNA diffusion coefficient when adding 250 nt chains instead of 125 nt is behind the augment of droplet viscosity at high RNA concentration shown in Fig. 5B. Moreover, when adding 10 nt chains, FUS (and RNA) diffusion considerably increases respect to those with 125 or 250 nt (at the same RNA concentration) due to the droplet density reduction (Fig. 5D). Interestingly, we also note that FUS, despite having the lower critical temperature to phase separate, and thus weaker LLPS-stabilising interactions than wt-TDP-43 and hnRNPA1 (Fig. 1B), displays the lowest protein diffusion of the set in absence of poly-U. Such intriguing fact, which might be related to patterning sequence effects^158^ or protein length^106^, highlights how beyond stability, condensate dynamics also entail intricate processes that need to be further investigated. In fact, methods promoting LLPS at lower protein concentration or enhancing protein mobility, such as by short RNA inclusion, could play therapeutic roles in preventing the emergence of pathological solid-like aggregates (by decreasing droplet density and viscosity) related to some neurodegenerative disorders such as amyotrophic lateral sclerosis or multisystem proteinopathy^15,50,62^.

## III. DISCUSSION AND CONCLUSIONS

Here, we investigate the dual effect of RNA in controlling the stability and dynamics of RNA-binding protein condensates. By means of a high-resolution sequence-dependent CG model for proteins and RNA^103,119,126^, we explore via MD simulations the underlying molecular and thermodynamic mechanisms enabling liquid-liquid phase separation of FUS, hnRNPA1 and TDP-43 along their corresponding prion-like and RRM domains, in presence *versus* absence of RNA poly-U strands. After validating the model by comparing the relative ability of the aforementioned proteins (without RNA) to phase separate against their experimental protein saturation concentration – finding a remarkable qualitative agreement between both simulations and experiments–, we characterize the condensates by determining their surface tension, key molecular contacts sustaining LLPS, and the protein conformational ensemble along the two phases. We find that highly inhomogeneous sequence contact maps, as those of wt-TDP-43, can lead to the emergence of largely heterogeneous droplets with low surface tension, where the exposure of PLD regions to the droplet interface deeply contributes in lowering *γ*, and favoring multidroplet emulsions^142,159,160^. However, such condensate heterogeneities can be significantly relieved when *α*–*α* helical PLD interactions are present, as recently hypothesized by Wang *et al.*^129^. Moreover, the analysis of the intermolecular contact maps within our droplets reveals the major importance of certain sequence domains of these RBPs in LLPS, such as the hnRNPA1 PLD-PLD interactions, or the FUS PLD-RGG interactions. Additionally, amino acid contacts such as G-G, R-Y, G-S, G-Y, K-F, or K-Y have been shown (Fig. S7-S9) to play a leading role in phase separation, highlighting the relevance of cation–*π* and electrostatic forces, besides hydrophobicity, in the physiological salt regime^61^. Also, the conformational protein ensemble inside the condensates has been demonstrated to be almost independent of temperature, in contrast to those measured in the diluted phase^86^. However, along the protein diluted-to-condensed transition, a significant variation to more extended conformational ensembles (to maximize protein molecular connectivity^116^) has been observed.

Our simulations with poly-U RNA also reveal how the formation of protein condensates is clearly enhanced at low poly-U concentration^50^, whereas inhibited at high poly-U/protein ratios^43,56,57^. The RNA concentration which promotes the highest increase in droplet stability is near the electroneutral charge point of poly-U/FUS and poly-U/hnRNPA1 mixtures (and also for both hnRNPA1 RRMs and A-LCD regions separately) in agreement with findings for LAF-1-PLD condensates^119^. We show how such boost in droplet stability is related to an increase of the condensate surface tension and liquid-network connectivity at low RNA ratios. In contrast, neither of the two studied TDP-43 variants, nor their RRMs together or individually, exhibited significantly LLPS enhancement through poly-U addition with this model. Besides, we demonstrate that beyond a certain strand length of ~100 nucleotides, the stability of the droplets for a given RNA concentration reaches a plateau, whereas below that minimum chain length, as for very short lengths (i.e., ~10 nt), it can even hinder phase separation^57^. These results have been shown to be related to the conformational structure and radius of gyration that RNA chains can adopt, which enable intermolecular binding between distinct proteins within the condensates. Overall, our results evidence how RBP condensate stability can be critically modulated by varying RNA concentration and length.

Finally, we focus on the transport properties of the RBP condensates as a function of RNA concentration and length. Our simulations demonstrate that while viscosity severely depends on the length of the added RNA chains – i.e., poly-U strands of 10 and 125 nt reduce droplet viscosity^43^, whereas 250-nucleotide strands moderately increase viscosity at high RNA concentration^49,58^ (Fig. 5) – protein diffusion hardly depends on poly-U length and mainly depends on droplet density, which in turn, is controlled by RNA concentration. The droplet viscosity gain with RNA length comes from the slower relaxation times and RNA diffusion in crowded environments by long RNA chains (Fig. 5 D). However, the addition of moderately short RNA strands (i.e., with a similar or slightly lower *R_g_* than those of the proteins) could help in promoting condensate dynamics without significantly destabilizing phase separation (Fig. 3 B). Our results suggest that the enhanced droplet dynamics at high RNA concentrations is mediated by a density reduction upon poly-U addition due to electrostatic repulsion. Taken together, our observations shed light on the crucial role of RNA (concentration and length) on the formation and phase behavior of RNA-protein complexes^54,58,125^. Moreover, the present work provides a novel estimation of the transport properties of protein condensates by means of computer simulations, which could pave the way for future studies characterizing the protein/RNA mobility in other relevant systems. Expanding our understanding on LLPS and the role of RNA in this process may drive solutions to precisely modulate aberrant liquid-to-solid transitions in the cell.

## IV. ACKNOWLEDGEMENTS

We acknowledge Dr R. Collepardo-Guevara and Dr J. A. Joseph for useful discussions. We also acknowledge the constructive comments from the reviewers for helping us to improve the manuscript. This project has received funding from the Oppenheimer Research Fellowship of the University of Cambridge. A. T. is funded by Universidad Politécnica de Madrid (ESTANCIAS-PIF-20-TYOSR8-13-4N0WPQ), A. G. is funded by an EPSRC studentship (EP/N509620/1) and a Winton scholarship. J. R. acknowledges funding from the Spanish Ministry of Economy and Competitivity (PID2019-105898GA-C22). J. R. E. also acknowledges funding from the Roger Ekins Research Fellowship of Emmanuel College. This work has been performed using resources provided by the Cambridge Tier-2 system operated by the University of Cambridge Research Computing Service (http://www.hpc.cam.ac.uk) funded by EP-SRC Tier-2 capital grant EP/P020259/1. The authors gratefully acknowledge the Universidad Politécnica de Madrid (www.upm.es) for providing computing resources on Magerit Supercomputer.

## V. AUTHOR CONTRIBUTIONS

J.R.E., and J.R. designed the research. A.R.T., and A.G. built the protein models, A.R.T. performed the simulations. A.R.T., J.R.E., and J.R. analyzed the data. A.R.T. and J.R.E. wrote the initial version of the manuscript. All authors contributed and edited the manuscript. J.R.E. and J.R. supervised the research.

## Notes

### Competing Interest Statement

The authors have declared no competing interest.

