## Supplementary Material for "RNA modulation of transport properties and stability in phase separated condensates"

(Dated: October 15, 2021)

* jorge.ramirez.upm.es

†

### SI. MODEL AND METHODS

#### A. Protein/RNA HPS model

In order to simulate the different studied proteins and RNA, we employ the LAMMPS Molecular Dynamics simulation package [1] with the recent reparameterization by Das *et al.* [2] of the chemically-accurate coarse-grained (CG) HPS protein model proposed by Dignon *et al.* [3]. For RNA, we use the new HPS-compatible CG model proposed by Regy *et al.* [4]. The coarse-grained model resolution, both for proteins and RNA, is of one bead per amino acid and nucleotide. In the model, the intrinsically disordered regions (IDRs) of the proteins are considered as fully flexible polymers, and the structured globular domains are treated as rigid bodies (where their conformations are taken from the Protein Data Bank (PDB) crystalline structure — see SII for the PBD codes) by using the rigid body integrator of LAMMPS [1]. Moreover, the interactions of the structured globular domains are scaled down by a 30% to account for the 'buried' amino acids as shown by Krainer *et al.* [5]. Also, in this model RNA strands are treated as flexible polymers.

The potential energy of the coarse-grained force field is given by:

$$E = E_{\text{Bonds}} + E_{\text{Electrostatic}} + E_{\text{Hydrophobic}} + E_{\text{Cation}-\pi}, \tag{S1}$$

where $E_{\text{Hydrophobic}}$, $E_{\text{Cation}-\pi}$ and $E_{\text{Electrostatic}}$ interactions are only applied between non-bonded beads and $E_{\text{Bonds}}$ between subsequent beads directly bonded to each other.

Bonded interactions between subsequent amino acid protein beads or consecutive RNA nucleotides are described by a harmonic potential:

$$E_{\text{Bonds}} = \sum_{\text{Protein/RNA bonds}} k(r - r_0)^2, \tag{S2}$$

where the equilibrium bond length is $r_0 = 5.0$Å between subsequent nucleotides and $r_0 = 3.81$Å between bonded amino acid beads. The spring constant is $k = 10\,\text{kJ}/(\text{mol}\text{Å}^2)$. Please see section SI C for further details on the value of this model parameter.

The electrostatic interactions, $E_{\text{Electrostatic}}$, among charged amino acids and RNA nucleotides are described by a Yukawa/Debye-Hückel potential of the form:

$$E_{\text{Electrostatic}} = \sum_i \sum_{j<i} \frac{1}{4\pi D} \frac{q_i q_j}{r} e^{-r/\kappa}, \tag{S3}$$

where $q_i$ and $q_j$ represent the charges of the beads $i$ and $j$ (amino acids or nucleotides), $D = 80$ is the dielectric constant of water, $r$ is the distance between the $i$th and $j$th beads, and $\kappa$= 1 $nm$ is the Debye screening length that mimics the implicit solvent (water and ions) at physiological salt concentration (~150mM of NaCl).

The hydrophobic interactions between different amino acid types and nucleotides are built upon a scale of amino acid and RNA nucleotide hydrophobicity based on a statistical potential derivation from contacts in PDB structures, and implemented through the functional form of an Ashbaugh/Hatch potential (see further details on these References [3, 4, 6]):

$$E_{\text{Hydrophobic}} = \sum_i \sum_{j<i} \begin{cases} 4\epsilon_{ij}\left[\left(\frac{\sigma_{ij}}{r}\right)^{12} - \left(\frac{\sigma_{ij}}{r}\right)^{6}\right] + (1-\lambda_{ij})\epsilon_{ij}, & r < 2^{1/6}\sigma_{ij} \\ \lambda_{ij} 4\epsilon_{ij}\left[\left(\frac{\sigma_{ij}}{r}\right)^{12} - \left(\frac{\sigma_{ij}}{r}\right)^{6}\right], & \text{otherwise,} \end{cases} \tag{S4}$$

where $\lambda_i$ and $\lambda_j$ are parameters that account for the hydrophobicity of the $i$th and $j$th interacting particles respectively, being $\lambda_{ij} = (\lambda_i + \lambda_j)/2$. The excluded volume of the different residues/nucleotides is given by $\sigma_i$ and $\sigma_j$, where $\sigma_{ij} = (\sigma_i + \sigma_j)/2$, and $r$ is the distance between the $ij$ particles. $\epsilon_{ij}$ (0.2 kcal/mol) is a fitting parameter to reproduce experimental single-IDR radius of gyration ($R_g$) [3]. When at least one of the $ij$ amino acids is part of a structured globular domain, $\epsilon_{ij}$ is scaled by a factor of 0.7 to account for 'buried' amino acids in globular domains. The concrete values for each amino acid and nucleotide $\sigma$, $q$, and $\lambda$ parameters can be found in References: Dignon *et al.* for proteins [3] and Regy *et al.* for RNA [4].

Finally, we consider an extra term for describing cation-$\pi$ interactions (only for the following set of pairs of amino acids (c-$\pi$:{Arg-Phe, Arg-Trp, Arg-Tyr, Lys-Phe, Lys-Trp and Lys-Tyr}):

$$E_{\text{cation-}\pi} = \sum_{i \in \text{c-}\pi} \sum_{j \in \text{c-}\pi \& j<i} 4\epsilon_{ij}\left[\left(\frac{\sigma_{ij}}{r}\right)^{12} - \left(\frac{\sigma_{ij}}{r}\right)^{6}\right], \tag{S5}$$

where $\sigma_{ij}$ is the same as in the hydrophobic interactions and $\epsilon_{ij}$ is $\epsilon_{ij} = 3.0 \text{kcal mol}^{-1}$ for all six cation-$\pi$ pairs as proposed in Ref. [2] (Approach 1). Consistently, the interaction of these amino acids is scaled down by a 30% when are found in structured globular domains.

### B. Simulation details

All simulations were carried out using LAMMPS [1] software. Direct Coexistence simulations (described in Section III) were carried out in the NVT ensemble using a Nosé-Hover thermostat [7] for the rigid bodies (crystalline structured domains of the proteins) and a Langevin thermostat [8] for the rest of the particles both with a relaxation time of 5 ps. The timestep for the Verlet integration of the equations of motion was chosen to be of 10 fs. NPT simulations for pure bulk protein liquids (see section V) were carried out at P = 1 bar using a Nosé-Hover barostat [1] and thermostat [7] with relaxation times of 50 ps and 5 ps respectively. For computational efficiency we use a cutoff of $3\sigma_{ij}$ for the cation-$\pi$ and hydrophobic interactions and 3.5 nm for the electrostatic ones [3]. We also turn off the interactions between particles that are part of the same globular structured domain (i.e., within the same rigid body).

### C. HPS model spring constant

The spring constant used in our simulations is $k = 10\,\mathrm{kJ/(mol\,\AA^2)}$, as described in the reparameterization of the HPS model with cation-$\pi$ interactions proposed by Das *et al* [2]. However, the HPS model was previously formulated with spring constants of $k = 10\,\mathrm{kJ/(mol\,\AA^2)}$ [3] and $k = 10\,\mathrm{kcal/(mol\,\AA^2)}$ [9]. To quantify the precise effect of $k$ in the phase behavior of IDRs, we perform Direct Coexistence simulations using the HPS model without cation-$\pi$ interactions to directly compare our results for different $k's$ with those provided by Dignon *et al.* for FUS40 [3] (Fig. S1).

We find that, despite the value of the spring constant does not dramatically change the phase diagram, the value that best reproduces the results by Dignon *et al.* [3] is $k = 10\,\mathrm{kJ/(mol\AA^2)}$. For this reason, and accordingly to the work of Das *et al.* in the HPS + cation-$\pi$ model [2], we use a spring constant of $k = 10\,\mathrm{kJ/(mol\AA^2)}$ or equivalently $k = 2.4\,\mathrm{kcal/(mol\AA^2)}$.

### D. Experimental validation of the HPS model without the cation–$\pi$ reparameterization

Here, we reproduce the comparison between the critical temperature measured in simula-

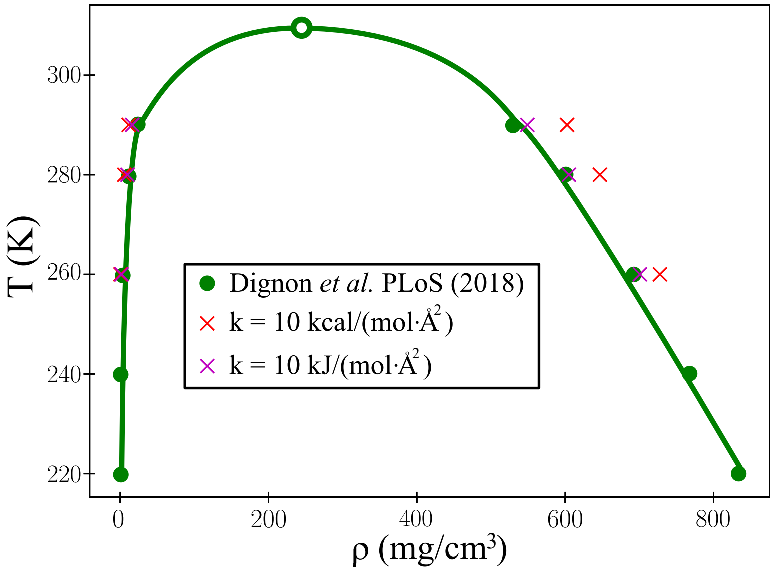


**FIG. S1:** Temperature-density phase diagram of FUS40 for two different spring constants ($k$) between bonded interactions, as indicated in the legend, see Eq. (S2). For comparison, we also include the phase diagram obtained by Dignon *et al* [3] (green circles). The empty circle shows the critical temperature obtained by Dignon *et al* in their original work ($T_c = 309.6K$). The solid line is included to help visualizing the phase diagram.

tions and the experimental protein saturation concentration, including results from the HPS model without the additional cation–$\pi$ interactions proposed by Das *et al.* [2]. We have run simulations of the HPS model for FUS, hnRNPA1, and wt-TDP-43, and we have computed their critical temperature using the law of rectilinear diameters and critical exponents [10]. The results for FUS-PLD can be directly adopted from the cation–$\pi$ reparameterization since the FUS-PLD sequence does not contain pairs of amino acids involved in cation–$\pi$ interactions. We show these results in Fig. S2.

Please recall that, although the absolute critical temperature of FUS-PLD is the same for both models, the critical temperature $T_c'$ of wt-TDP-43, used to normalize the data of each set, is not. As it can be seen in Figure S2, the correlation between the critical temperature from simulations and the experimental saturation concentration of the proteins exhibited by the HPS model+cation–$\pi$ (Figure 1 of the main text), is no longer present in the case of the HPS model. More notably, the critical temperature of FUS is way below the critical point of FUS-PLD, which clearly contradicts the experimental trend. The saturation concentration of FUS-PLD is approximately one order of magnitude greater than the corresponding full-sequenced FUS, and thus, the critical point of the full protein is expected to be greater than the $T_c$ of FUS-PLD. These results show the necessity of including an extra term for cation–$\pi$

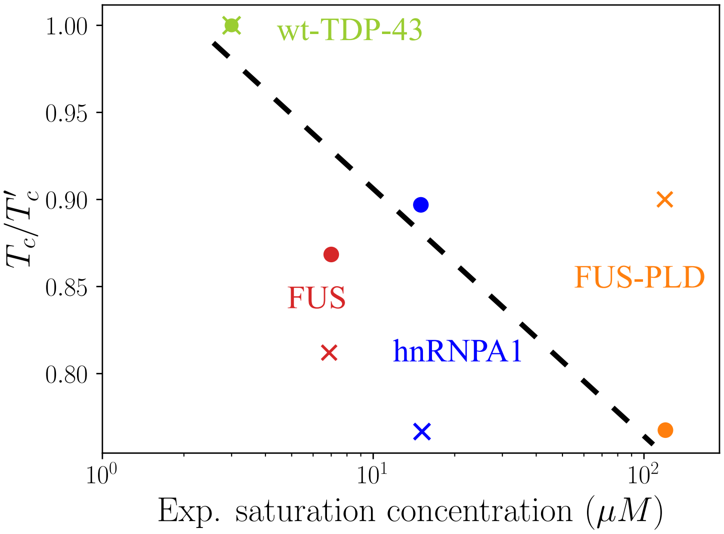


**FIG. S2:** Comparison of the renormalized critical temperature measured in simulations with the HPS+cation–$\pi$ (filled circles) and the HPS models (crosses) against the experimental saturation concentration of the proteins. The experimental saturation concentrations at physiological salt conditions were obtained from the references provides in the main text. Temperature is renormalized by the highest critical temperature $T'_c$ of each set, corresponding to wt-TDP-43 in both cases.

interactions in the HPS model as proposed by Das *et al.* [2].

### SII. SEQUENCES AND PDBS OF THE PROTEINS USED

#### FUS

MASNDYTQQATQSYGAYPTQPGQGYSQQSSQPYGQQSYSGYSQSTDTSGYGQSSYSSYGQSQNTG
YGTQSTPQGYGSTGGYGSSQSSQSSYGQQSSYPGYGQQPAPSSTSGSYGSSSQSSSYGQPQSGSYSQ
QPSYGGQQQSYGQQQSYNPPQGYGQQNQYNSSSGGGGGGGGGNYGQDQSSMSSGGGSGGGYG
NQDQSGGGGSGGYGQQDRGGRGRGGSGGGGGGGGGGYNRSSGGYEPRGRGGGRGGRGGMGGS
DRGGFNKFGGPRDQGSRHDSEQDNSDNNTIFVQGLGENVTIESVADYFKQIGIIKTNKKTGQPMIN
LYTDRETGKLKGEATVSFDDPPSAKAAIDWFDGKEFSGNPIKVSFATRRADFNRGGGNGRGGRGR
GGPMGRGGYGGGGSGGGGRGGFPSGGGGGGGQQRAGDWKCPNPTCENMNFSWRNECNQCKA
PKPDGPGGGPGGSHMGGNYGDDRRGGRGGYDRGGYRGRGGDRGGFRGGRGGGDRGGFGPGK
MDSRGEHRQDRRERPY

#### FUS-PLD

MASNDYTQQATQSYGAYPTQPGQGYSQQSSQPYGQQSYSGYSQSTDTSGYGQSSYSSYGQSQNTG
YGTQSTPQGYGSTGGYGSSQSSQSSYGQQSSYPGYGQQPAPSSTSGSYGSSSQSSSYGQPQSGSYSQ
QPSYGGQQQSYGQQQSYNPPQGYGQQNQYNS

#### hnRNPA1

MSKSESPKEPEQLRKLFIGGLSFETTDESLRSHFEQWGTLTDCVVMRDPNTKRSRGFGFVTYATVE
EVDAAMNARPHKVDGRVVEPKRAVSREDSQRPGAHLTVKKIFVGGIKEDTEEHHLRDYFEQYGKI
EVIEIMTDRGSGKKRGFAFVTFDDHDSVDKIVIQKYHTVNGHNCEVRKALSKQEMASASSSQRGRS
GSGNFGGGRGGGFGGNDNFGRGGNFSGRGGFGGSRGGGGYGGSGDGYNGFGNDGGYGGGGPG
YSGGSRGYGSGGQGYGNQGSGYGGSGSYDSYNNGGGGGFGGGSGSNFGGGGSYNDFGNYNNQSS
NFGPMKGGNFGGRSSGPYGGGGQYFAKPRNQGGYGGSSSSSSYGSGRRF

#### hnRNPA1-PLD

GDGYNGFGNDGGYGGGGPGYSGGSRGYGSGGQGYGNQGSGYGGSGSYDSYNNGGGGGFGGGSG
SNFGGGGSYNDFGNYNNQSSNFGPMKGGNFGGRSSGPYGGGGQYFAKPRNQGGYGGSSSSSSYGS
GRRF

**hnRNPA1-RRMs**

MSKSESPKEPEQLRKLFIGGLSFETTDESLRSHFEQWGTLTDCVVMRDPNTKRSRGFGFVTYATVE
EVDAAMNARPHKVDGRVVEPKRAVSREDSQRPGAHLTVKKIFVGGIKEDTEEHHLRDYFEQYGKI
EVIEIMTDRGSGKKRGFAFVTFDDHDSVDKIVIQKYHTVNGHNCEVRKALSKQ

**hnRNPA1-A-LCD**

MASASSSQRGRSGSGNFGGGRGGGFGGNDNFGRGGNFSGRGGFGGSRGGGGYGGSGDGYNGFG
NDGSNFGGGGSYNDFGNYNNQSSNFGPMKGGNFGGRSSGPYGGGGQYFAKPRNQGGYGGSSSSSS
YGSGRRF

**TDP-43**

MSEYIRVTEDENDEPIEIPSEDDGTVLLSTVTAQFPGACGLRYRNPVSQCMRGVRLVEGILHAPDAG
WGNLVYVVNYPKDNKRKMDETDASSAVKVKRAVQKTSDLIVLGLPWKTTEQDLKEYFSTFGEVL
MVQVKKDLKTGHSKGFGFVRFTEYETQVKVMSQRHMIDGRWCDCKLPNSKQSQDEPLRSRKVFV
GRCTEDMTEDELREFFSQYGDVMDVFIPKPFRAFAFVTFADDQIAQSLCGEDLIIKGISVHISNAEPK
HNSNRQLERSGRFGGNPGGFGNQGGFGNSRGGGAGLGNNQGSNMGGGMNFGAFSINPAMMAAA
QAALQSSWGMMGMLASQQNQSGPSGNNQNQGNMQREPNQAFGSGNNSYSGSNSGAAIGWGSASN
AGSGSGFNGGFGSSMDSKSSGWGMMSEYIRVTEDENDEPIEIPSEDDGTVLLSTVTAQFPGACGLR
YRNPVSQCMRGVRLVEGILHAPDAGWGNLVYVVNYPKDNKRKMDETDASSAVKVKRAVQKTSD
LIVLGLPWKTTEQDLKEYFSTFGEVLMVQVKKDLKTGHSKGFGFVRFTEYETQVKVMSQRHMID
GRWCDCKLPNSKQSQDEPLRSRKVFVGRCTEDMTEDELREFFSQYGDVMDVFIPKPFRAFAFVTF
ADDQIAQSLCGEDLIIKGISVHISNAEPKHNSNRQLERSGRFGGNPGGFGNQGGFGNSRGGGAGLGN
NQGSNMGGGMNFGAFSINPAMMAAAQAALQSSWGMMGMLASQQNQSGPSGNNQNQGNMQREP
NQAFGSGNNSYSGSNSGAAIGWGSASNAGSGSGFNGGFGSSMDSKSSGWGM

**TDP-43-PLD**

GRFGGNPGGFGNQGGFGNSRGGGAGLGNNQGSNMGGGMNFGAFSINPAMMAAAQAALQSSWG
MMGMLASQQNQSGPSGNNQNQGNMQREPNQAFGSGNNSYSGSNSGAAIGWGSASNAGSGSGFNG
GFGSSMDSKSSGWGM

The following Protein Data Bank (PDB) codes were used to build the globular structured domains of: FUS (residues from 285–371 (PDB code: 2LCW) and from 422–453 (PDB code:

6G99)), wt-TDP-43 (residues 2-38, 40-49 and 51-79 all included in the same PDB, (PDB code: 5MDI) and from residues 193-267 (PDB code: 1WF0)), h-TDP-43 (additionally to the structured domains of wt-TDP-43, this variant has an $\alpha$–helical from residues 307-349 (PBD code: 2N2C)) and hnRNPA1 (residues from 8-91 and 103-181 in the same PDB (PDB code: 1L3K)). The intrinsically disordered regions not included in the PDBs were built using the VMD software [11].

### SIII. PHASE DIAGRAM CALCULATIONS VIA DIRECT COEXISTENCE SIMULATIONS

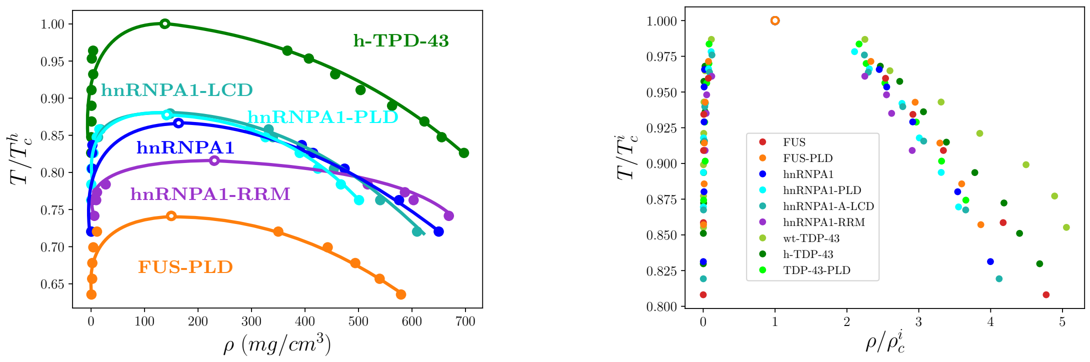


**(a)** **(b)**

**FIG. S3:** **(a)** Temperature-density phase diagram (normalised by $T_c^h$, where $T_c^h$ is the critical temperature of h-TDP-43, $T_c^h$=472K) for FUS-PLD (orange), hnRNPA1-RRM (purple), hnRNPA1-A-LCD (turquoise), hnRNPA1-PLD (cyan) and h-TDP-43 (green). The critical density and temperature (depicted by empty circles) have been obtained by using the law of rectilinear diameters and critical exponents, Eqs. (S6) and (S7). **(b)** Temperature-density phase diagram for all the studied proteins where both temperature and density are renormalized by their own critical temperature ($T_c^i$) and critical density ($\rho_c^i$). The shown sequences are: FUS (red), hnRNPA1 (dark blue), wt-TDP-43 (lime green), h-TDP-43 (dark green), TDP-43-PLD (light green), hnRNPA1-A-LCD (turquoise), hnRNPA1-PLD (cyan), FUS-PLD (orange) and hnRNPA1-RRM (purple). Both critical density and temperature for each protein were obtained as described in (a).

| Protein | Chains ($N_c$) | Protein residues |
|---|---|---|
| FUS | 24 | 526 |
| FUS-PLD | 80 | 163 |
| hnRNPA1 | 40 | 372 |
| hnRNPA1-PLD | 100 | 132 |
| hnRNPA1-RRM | 80 | 184 |
| hnRNPA1-A-LCD | 200 | 135 |
| TDP-43 | 32 | 414 |
| TDP-43-PLD | 100 | 141 |

**TABLE S1:** Employed systems sizes in the Direct Coexistence simulations of each protein type (in pure component) including the number of protein replicas ($N_c$) and the number of amino acids per protein. Simulations including poly-U contain the same number of proteins, except hnRNPA1-A-LCD, which is reduced to $N_c = 100$ to keep similar RNA/protein ratios in all systems.

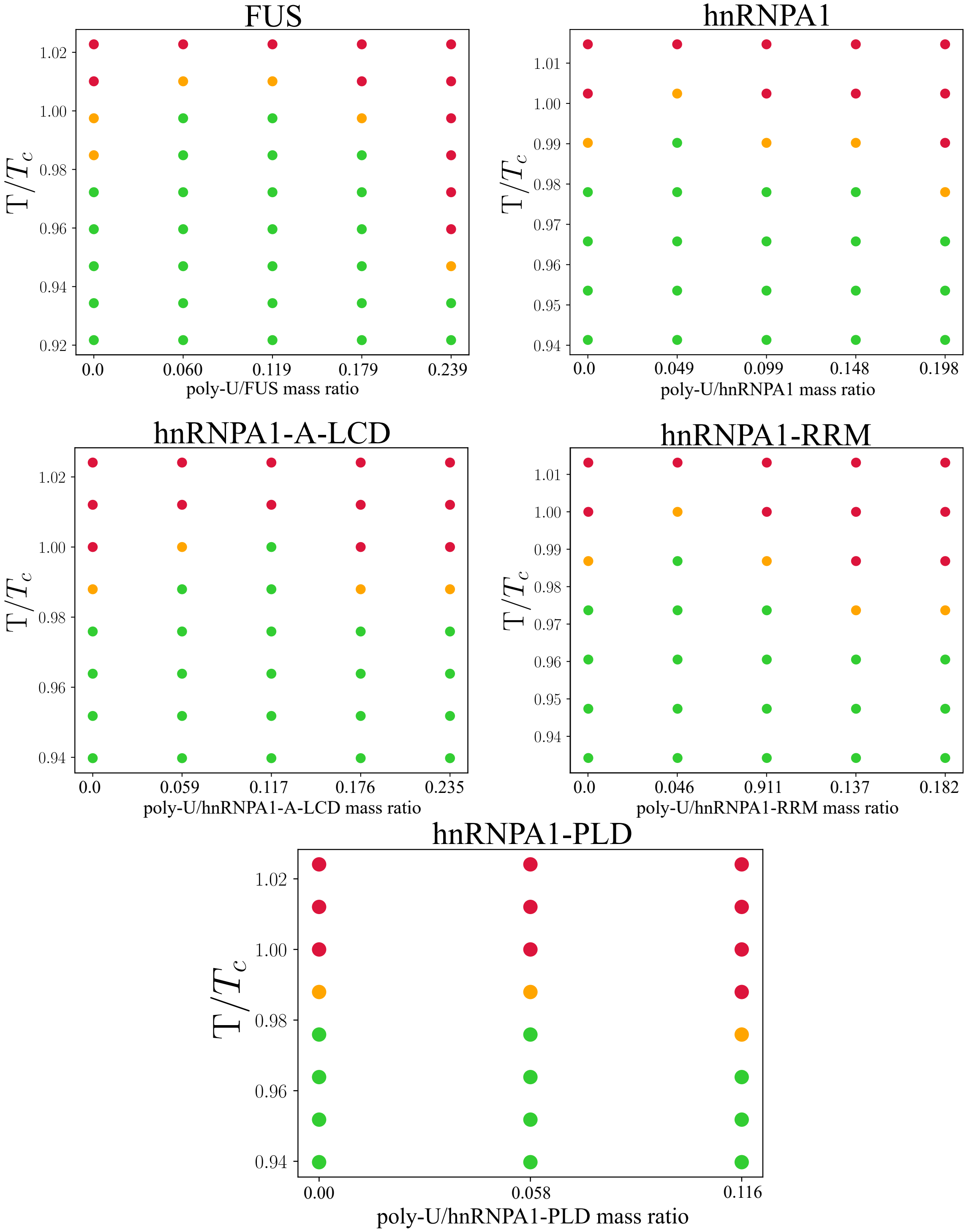


**FIG. S4:** Phase behaviour of FUS, hnRNPA1, hnRNPA1-A-LCD, hnRNPA1-RRM and hnRNPA1-PLD condensates at different poly-U concentrations as a function of temperature (renormalized by the critical temperature ($T_c$) in absence of poly-U (Table S2). Green circles indicate temperatures/concentrations where phase separation was observed in our DC simulations, and red circles where no phase separation was observed. Orange dots show the limit between both regimes. Note that LLPS cannot be directly observed by means of DC simulations just below the critical temperature due to finite size effects, and thus, the highest temperature at which we can usually observe LLPS by means of DC simulations is $T/T_c \sim 0.98$.

We perform Direct Coexistence (DC) simulations [12–15] to compute the phase diagram of the different proteins and protein/RNA mixtures (See Table S1 for the employed system sizes). Within DC simulations, the two coexisting phases of the system are simulated in the same simulation box. In our case, we place a high-density protein liquid and a very low-density one. We employ a rectangular box, with an elongated side perpendicular to the interfaces (long enough to capture the bulk density of each phase), while the parallel sides are chosen such that proteins and RNA cannot interact with themselves along the periodic boundary conditions. We run NVT simulations until equilibrium is reached. Then, we measure the equilibrium coexisting densities of both phases along the long side of the box, excluding the fluctuations of the interfaces and keeping the center of mass of the slab fixed. We repeat this procedure at different temperatures until we reach the critical temperature. To avoid finite system-size effects close to the critical point we evaluate the critical temperature ($T_c$) and density ($\rho_c$) using the law of critical exponents and rectilinear diameters [10]:

$$(\rho_l - \rho_v)^{\alpha} = s_1(1 - \frac{T}{T_c}) \tag{S6}$$

$$\frac{\rho_l + \rho_v}{2} = \rho_c + s_2(T_c - T), \tag{S7}$$

where $\rho_l$ and $\rho_v$ refer to the densities of the condensed and the diluted phases respectively, $s_1$ and $s_2$ are fitting parameters, and $\alpha = 3.06$ accounts for the critical exponent of the three dimensional Ising model [10].

| Protein | Method 1 ($T_c/K$) | Method 2 ($T_c/K$) |
|---|---|---|
| FUS | 396 | 400 |
| FUS-PLD | 350 | 347 |
| hnRNPA1 | 409 | 407 |
| hnRNPA1-RRM | 385 | 390 |
| hnRNPA1-PLD | 414 | 411 |
| hnRNPA1-A-LCD | 415 | 413 |
| h-TDP-43 | 472 | 468 |
| wt-TDP-43 | 456 | 448 |
| TDP-43-PLD | 366 | 367 |

**TABLE S2:** Comparison of the critical temperatures estimated by using the law of rectilinear diameters and critical exponents (Method 1), and by fitting the surface tension data as a function of temperature using the fit given in Eq. (S9) (Method 2).

| poly-U / Protein | 0 ($T_c/K$) | 1 ($T_c/K$) | 2 ($T_c/K$) | 3 ($T_c/K$) | 4 ($T_c/K$) |
|---|---|---|---|---|---|
| FUS | 396 | 405 | 401 | 398 | 386 |
| hnRNPA1 | 409 | 412 | 417 | 407 | 405 |
| hnRNPA1-A-LCD | 415 | 418 | 419 | 411 | 409 |
| hnRNPA1-PLD | 414 | 416 | 406 | - | - |
| hnRNPA1-RRM | 385 | 391 | 382 | - | - |

**TABLE S3:** Comparison of the estimated critical temperatures (using Method 1) for the different poly-U/protein mixtures as a function of poly-U concentration, given in number of added poly-U strands of 250 nucleotides. The number of proteins in each system is that given in Table S1, and is constant for all concentrations. The equivalence in poly-U/protein mass ratio with the number of added of 250 nt poly-U strands can be extracted by comparison with Fig. S4.

### SIV. INTERFACIAL FREE ENERGY

As previously explained, in DC simulations two phases coexist in the same simulation box. Since one of the box sides (*e.g.* z-axis) is longer than the other two, the bulk density of the two phases can be conveniently measured along the corresponding axis. Moreover, when the system is equilibrated, the surface tension can be also evaluated from the computed pressure tensor. By means of the following expression, the interfacial free energy ($\gamma$) can be evaluated [16]:

$$\gamma = \frac{L_N}{2}(p_N - p_T), \tag{S8}$$

where $p_N$ denotes the normal component of the pressure tensor perpendicular to the interface, $p_T$ represents the average of the tangential components of the pressure tensor, $L_N$ denotes the length of the long side of the simulation box and the 2 factor accounts for the presence of two interfaces in the simulation box. The surface tensions shown in Fig. S5 (and Figs. 2(A) and 3(F) of the main text) have been computed using this expression. Furthermore, $\gamma$ values can be also used to alternatively estimate the critical temperature by assuming the following scaling [10]:

$$\gamma = A(T_c - T)^{1.26}, \tag{S9}$$

where $T_c$ and $A$ are fitting parameters. The critical temperature can be estimated as the temperature at which $\gamma$ becomes zero [17].

### SV. CONFORMATIONAL PROTEIN ENSEMBLE

We analyse the protein conformational ensemble of the different sequences along the condensed and diluted liquid phases. For this purpose, we compute the histogram of the radius of gyration ($R_g$) distribution function of the proteins in both phases. To measure $R_g$ in the diluted phase, we run NVT simulations with a single protein at different temperatures and at the coexisting density of each temperature according to the phase diagram. Upon reaching equilibrium, we compute the $R_g$ histograms along the simulation. To measure $R_g$ inside the condensates, we first equilibrate the liquid of the given protein at the desired temperature and density, and then, we run NVT simulations in which we compute the $R_g$ of all the protein replicas along time. The averaged $R_g$ distributions over time and protein replicas are represented in Fig. S6 for the different studied sequences within the condensates (continuous lines) and in the diluted phase (dashed ones) for the lowest and highest temperature at which phase separation was observed.

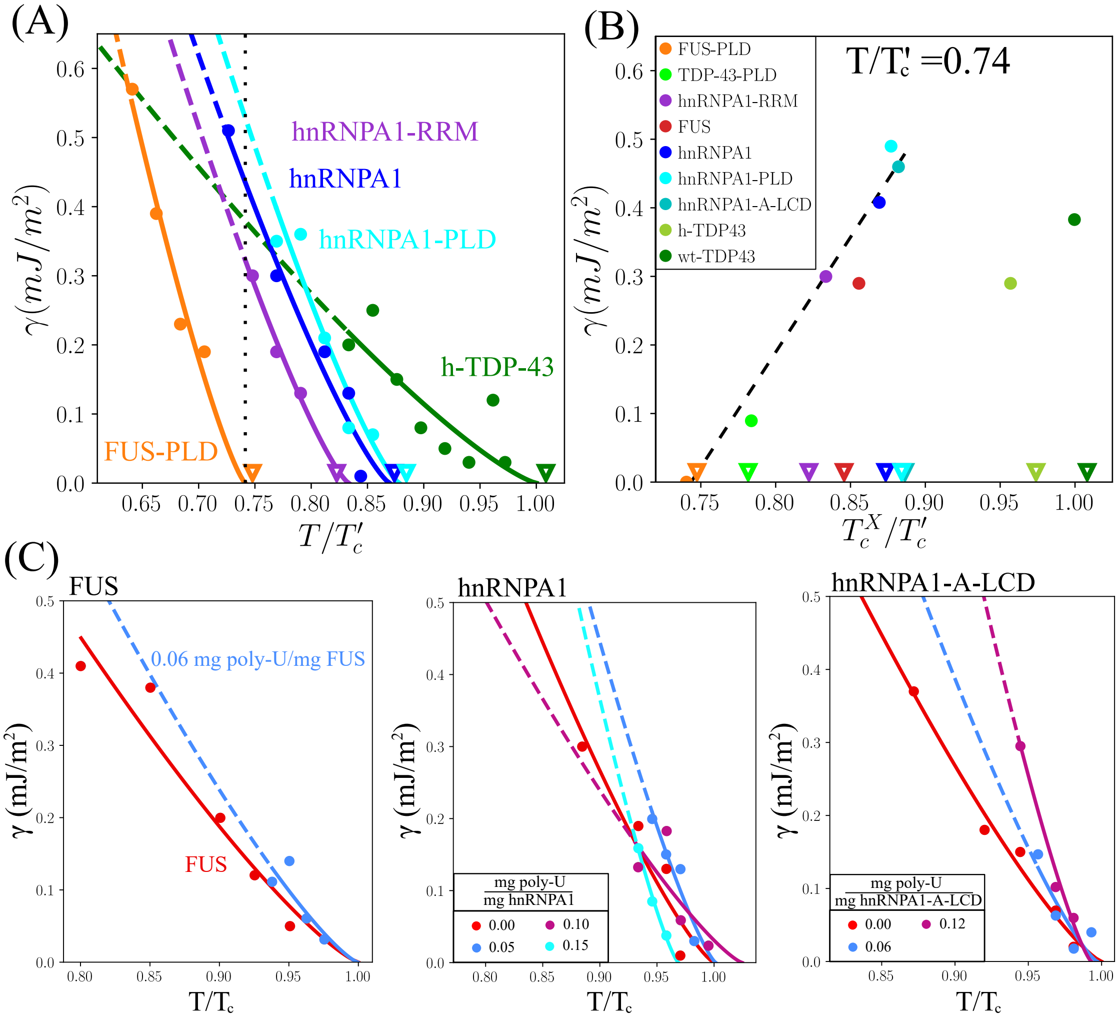


**FIG. S5:** (A): Droplet surface tension ($\gamma$) of hnRNPA1-RRM, hnRNPA1, hnRNPA1-PLD, h-TDP-43 and FUS-PLD condensates as a function of $T/T_c'$ where $T_c'$ is the critical temperature of h-TDP-43 as obtained by the extrapolation method explained above. Filled circles indicate the value of $\gamma$ as obtained from DC simulations and solid curves are the representation of the curves recovered by fitting our data to Eq. (S9). Dashed curves depict the predicted surface tension at low T as extrapolated from the fit. Empty triangles represent the predicted (renormalized by $T_c'$) critical point of each system ($T_c^X$) using the laws of rectilinear diameters and critical exponents. See table S2 for further details. Black dotted line indicates the temperature selected for panel (B). (B) Interfacial tension measured at $T/T_c' = 0.74$ (see panel (A)) for all the systems as a function of the critical temperature obtained from DC simulations using the fit of data to Eq. (S9). The critical temperature of each system ($T_c^X$) is normalized by $T_c'$ which corresponds to the critical temperature of h-TDP-43. Empty triangles show the critical temperatures (normalized by $T_c'$) using the law of rectilinear diameters and critical exponents. The black dashed line shows the trend in $\gamma$ *vs.* $T_c^X/T_c$ which works for all proteins except for both variants of TDP-43 as expected. (C): Surface tension of the poly-U-protein condensates for FUS, hnRNPA1 and hnRNPA1-A-LCD. Filled circles show the data obtained from simulations and solid lines denote scaling fit given in Eq. (S9). The results for the surface tension of the RNA-protein mixtures are more unstable due to the proximity to the critical point and thus, the estimation of $T_c$ is less accurate. Dashed curves represent the extrapolation of the interfacial tension at low temperatures.

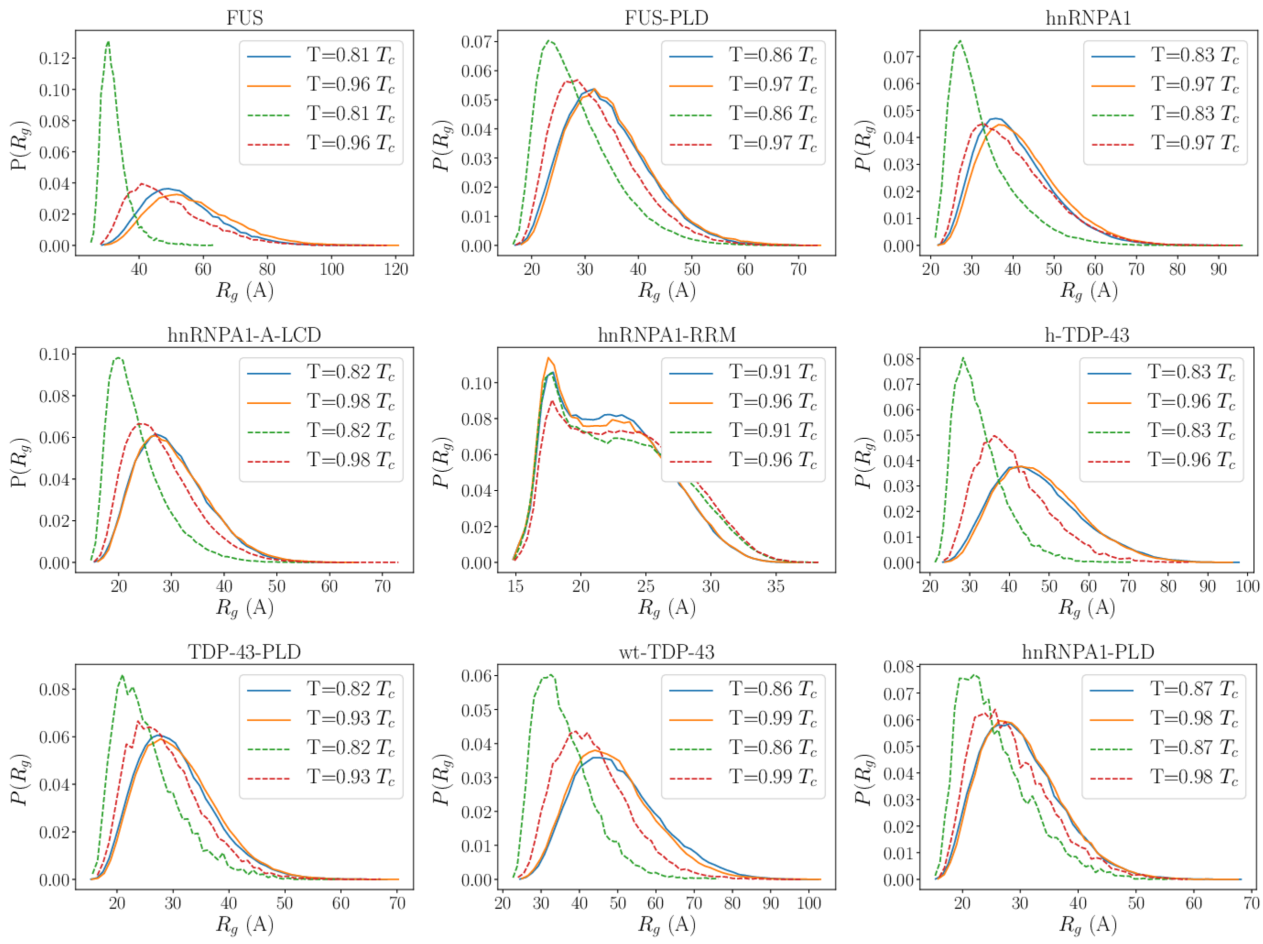


**FIG. S6:** Normalised radius of gyration histograms of the different studied proteins within the condensate (solid curves) and within the diluted phase (dashed lines) at the temperatures indicated in the legend. Note that $T_c$ refers to the critical temperature of each system in pure component.

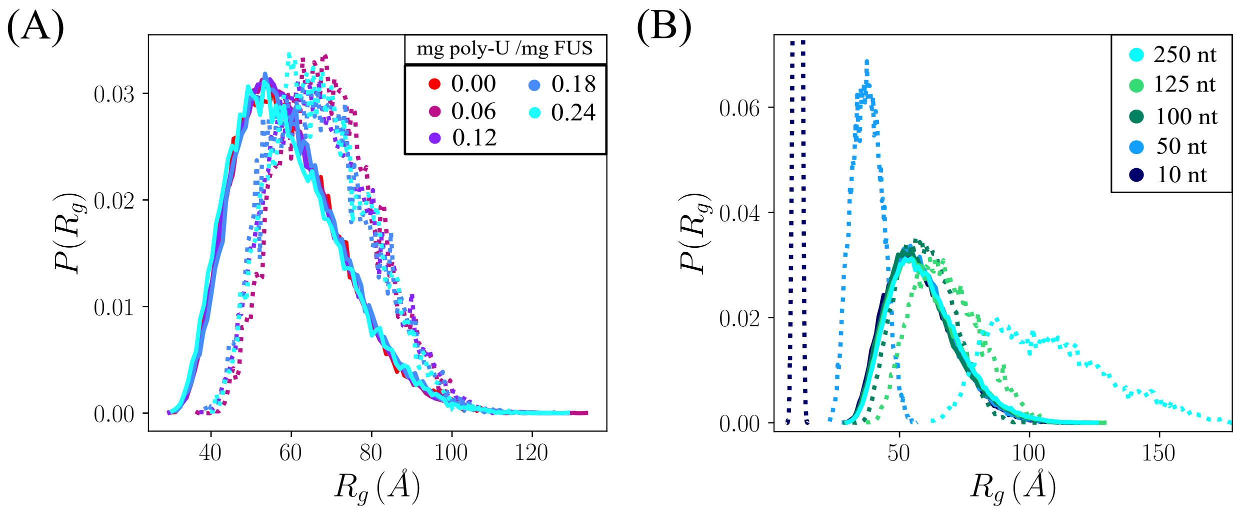


**FIG. S7:** (A) Normalised radius of gyration histograms of FUS (solid lines) and poly-U strands (dashed lines) within the condensate for different RNA-FUS ratios (as indicated in the legend) at $T \approx 0.97T_c$, where $T_c$ refers to the critical temperature of FUS in absence of RNA. (B) Normalised radius of gyration histograms of FUS (solid lines) and poly-U strands (dashed lines) within the condensate for different poly-U strand lengths and for a constant poly-U/FUS mass ratio of 0.06. The temperature of the studied systems is $T \approx 0.97T_c$, where $T_c$ is the critical temperature in absence of RNA.

### SVI. PROTEIN/RNA CONTACT MAPS

The intermolecular contact maps between proteins inside the condensates are computed from DC trajectories. For all systems, the contacts were computed at $T/T_c \approx 0.9$ in absence of poly-U and at $T/T_c \approx 0.95$ in presence of poly-U, being $T_c$ the critical temperature of each corresponding system (when poly-U is present, $T_c$ refers to the critical temperature of the mixture at the considered poly-U concentration). Typically, molecular contacts are tracked through a distance criterion, and it is normally assumed that the contact map relative frequency (not absolute frequency) is in general independent of the chosen cut-off distance (for reasonable cut-off values) used in such calculations. However, in order to accurately compute the most relevant and frequent residue-residue contact pairs enabling LLPS, it is essential to specifically consider the actual parametrization of each amino acid in terms of excluded volume and minimum potential energy interacting distance. For that reason, we used a 'smart' sequence dependent cut-off distance equal to $1.2\sigma_{ij}$, where $\sigma_{ij}$ accounts for the mean excluded volume of the specific $i$th and $j$th amino acids. Since the minimum of the employed potential is located at $2^{1/6}\sigma_{ij} \approx 1.122\sigma_{ij}$, we set our cut-off distance slightly beyond that point, at $1.2\sigma_{ij}$. Using this novel sequence-dependent cut-off scheme for each amino acid pair interaction, we can better exclude neighboring contacts that are coincidentally close to real interacting amino acids along the sequence that indeed are positively contributing to sustain LLPS.

In Figs. S8 and S12 we show the protein contact maps (averaged over all the equilibrium configurations and protein replicas) in absence (S8) *versus* presence of poly-U (at the poly-U/protein mass ratio that maximises droplet stability, see caption of Fig. S12 for the specific poly-U/protein mass fraction of each system). Given that the poly-U sequence is only composed by uridines, we also include side bars in the corresponding maps of Fig. S12, which show all the contacts between uridines and the given protein amino acids. In Fig. S12 (B), we plot the total number of intermolecular contacts between FUS (top) and hnRNPA1 (middle) domains with poly-U averaged over all the equilibrium configurations and protein regions as labelled in the figure.

Moreover, the ten most frequent intermolecular protein contacts within the condensates (according to the force field [2, 3]) are provided in Figures S9-S11 for pure component droplets, and in Fig. S13 for droplets with poly-U. These figures include the top ten most repeated contacts evaluated with the sequence-dependent cut-off described above (panel (A)), and the ten most frequent contacts after renormalization by the relative abundance of each amino acid along the protein sequence (panel (B)). Furthermore, in panel (C), we provide the natural abundance of the different amino acids along each protein sequence. The top ten contacts provided in panel (A) indicates the number of intermolecular contacts between pairs of amino acids per protein and configuration (averaged over all of them). That magnitude over the number of amino acids in the protein sequence that are involved in every pairwise contact is shown in B. The same information but for the poly-U/protein mixtures described in Fig. S12 is provided in Fig. S13.

Finally, in Figure S14, we provide information of the number of contacts between FUS-FUS, FUS–poly-U and the total number of contacts of different FUS-RNA mixtures. In panel (A), the number of contacts are calculated for different poly-U concentrations and strand lengths of 125 nt. On the other hand, panel (B) shows the same analysis as a function of the strand length for 3 different cases (10 nt, 125 nt, and 250 nt) and at a constant concentration of 0.06 mg poly-U/mg FUS. All results have been obtained from the same simulations employed to compute viscosity and droplet diffusion.

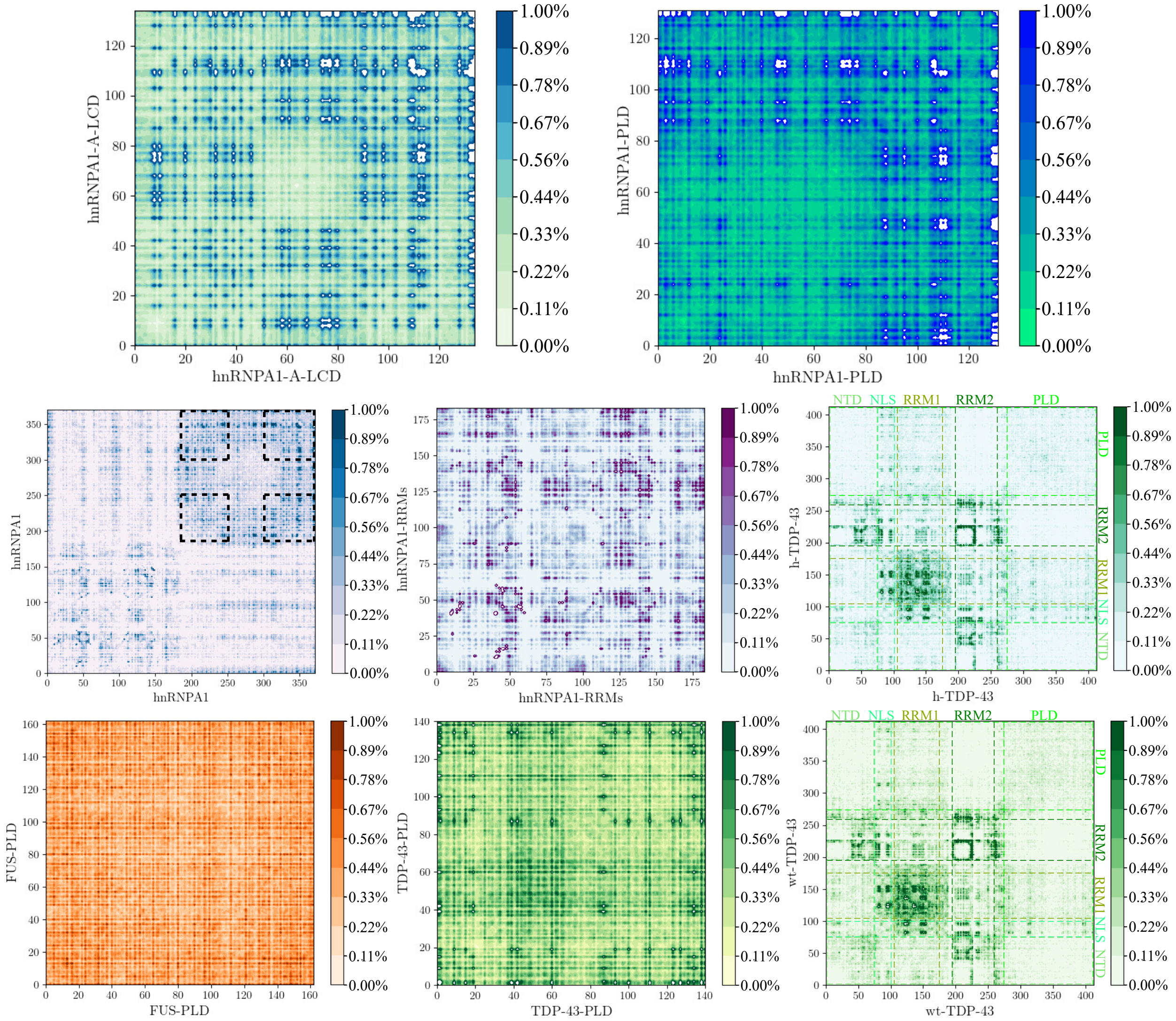


**FIG. S8:** Average number of intermolecular protein contacts per protein replica in percentage (where 100% would mean that all protein replicas in the condensate have a given contact in all configurations) for hnRNPA1-A-LCD, hnRNPA1-PLD, hnRNPA1, hnRNPA1-RRMs, FUS-PLD, TDP-43-PLD, wt-TDP-43 and h-TDP-43 condensates in pure component measured at $T/T_c = 0.95$, being $T_c$ the corresponding critical temperature of each protein type (see Table S2). Dashed squares in the contact map of hnRNPA1 show the two corresponding sequences of hnRNPA1-A-LCD contained in hnRNPA1. The different protein domains in wt-TDP-43 and h-TDP-43 are indicated by dashed lines.

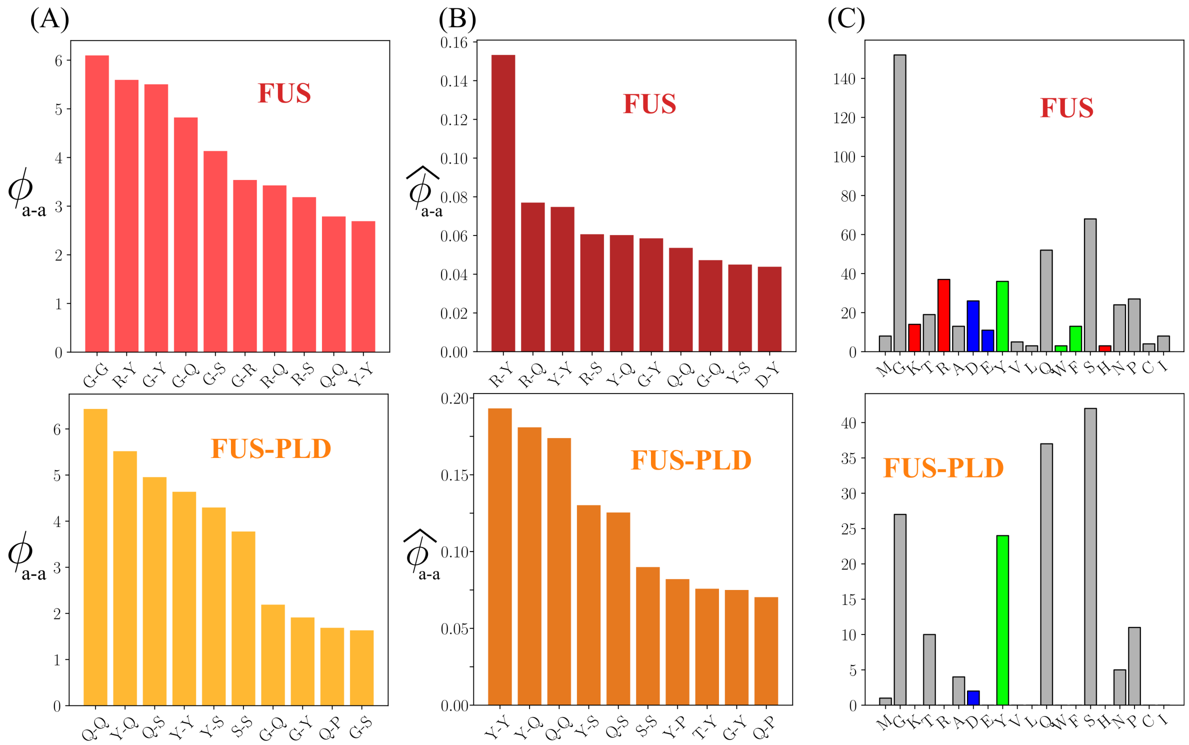


**FIG. S9:** (A) Average number of the ten most repeated intermolecular amino acid contacts per protein ($\phi_{a-a}$) within FUS and FUS-PLD droplets (in absence of poly-U) at $T/T_c = 0.9$ (see Table S2 for the critical temperature $T_c$ of each protein). (B) The same as in (A) but renormalized by the protein amino acid abundance ($\hat{\phi_{a-a}}$). Note that for every given pairwise contact interaction, we renormalize by the sum of the amount of the two amino acids involved in the contact interaction divided by two. If the amino acid pair is between same type of amino acids, we then recover the natural abundance of that amino acid type in the sequence. (C) Abundance of each amino acid type in FUS and FUS-PLD sequences. The color code indicates positively charged (red), negatively charged (blue) and aromatic residues (green), while the rest of amino acids are labelled in grey.

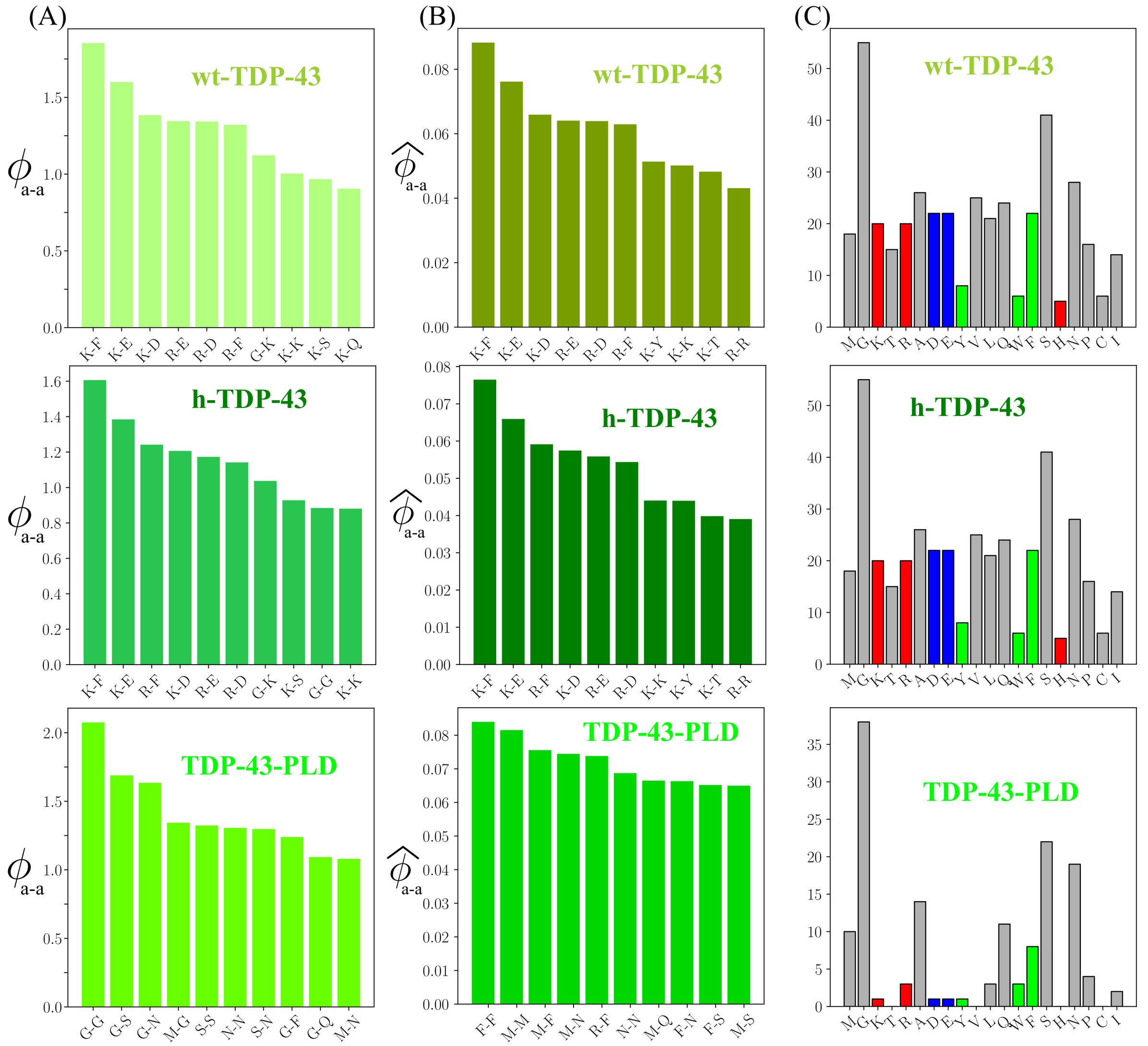


**FIG. S10:** (A) Average number of the ten most repeated intermolecular amino acid contacts per protein ($\phi_{a-a}$) within wt-TDP-43, h-TDP-43 and TDP-43-PLD droplets (in absence of poly-U) at $T/T_c = 0.9$ (see Table S2 for the critical temperature $T_c$ of each protein). (B) The same as in (A) but renormalized by the protein amino acid abundance ($\hat{\phi}_{a-a}$). Note that for every given pairwise contact interaction, we renormalize by the sum of the amount of the two amino acids involved in the contact interaction divided by two. If the amino acid pair is between the same type of amino acids, we then recover the natural abundance of that amino acid type in the sequence. (C) Abundance of each amino acid type in the sequences of wt-TDP-43, h-TDP-43 and TDP-43-PLD. The color code indicates positively charged (red), negatively charged (blue) and aromatic residues (green), while the rest are labelled in grey.

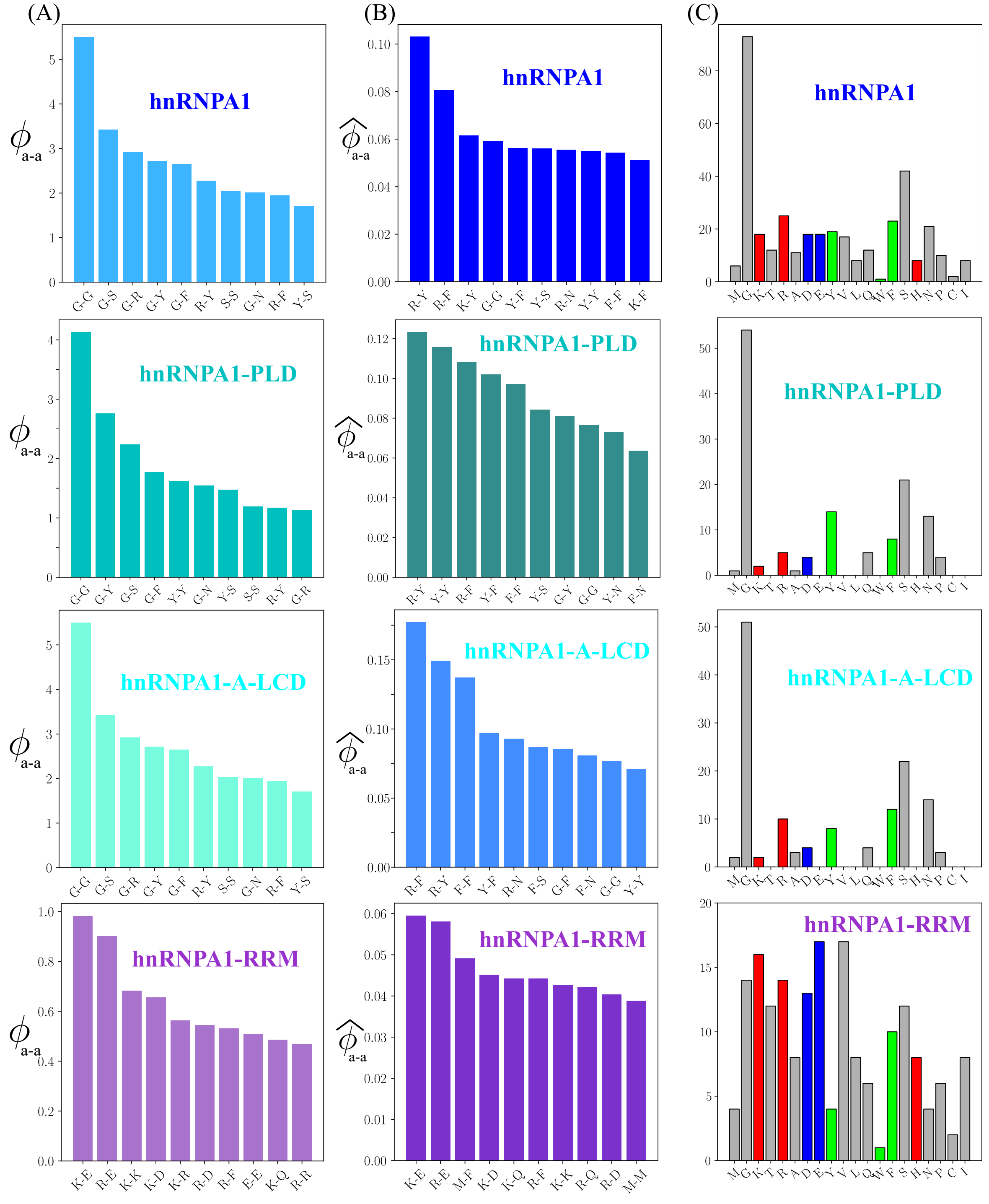


**FIG. S11:** (A) Average number of the ten most repeated intermolecular amino acid contacts per protein ($\phi_{a-a}$) within hnRNPA1, hnRNPA1-PLD, hnRNPA1-A-LCD and hnRNPA1-RRM droplets (in absence of poly-U) at $T/T_c = 0.9$ (see Table S2 for the critical temperature of each protein). (B) The same as in (A) but renormalized by the protein amino acid abundance ($\hat{\phi}_{a-a}$). Note that for every given pairwise contact interaction, we renormalize by the sum of the amount of the two amino acids involved in the contact pair divided by two. If the amino acid pair is between the same type of amino acids, we then recover the natural abundance of that amino acid type in the sequence. (C) Abundance of each amino acid type in the sequences of hnRNPA1, hnRNPA1-PLD, hnRNPA1-RRM and hnRNPA1-A-LCD proteins. The color code indicates positively charged (red), negatively charged (blue) and aromatic residues (green), while the rest are labelled in grey.

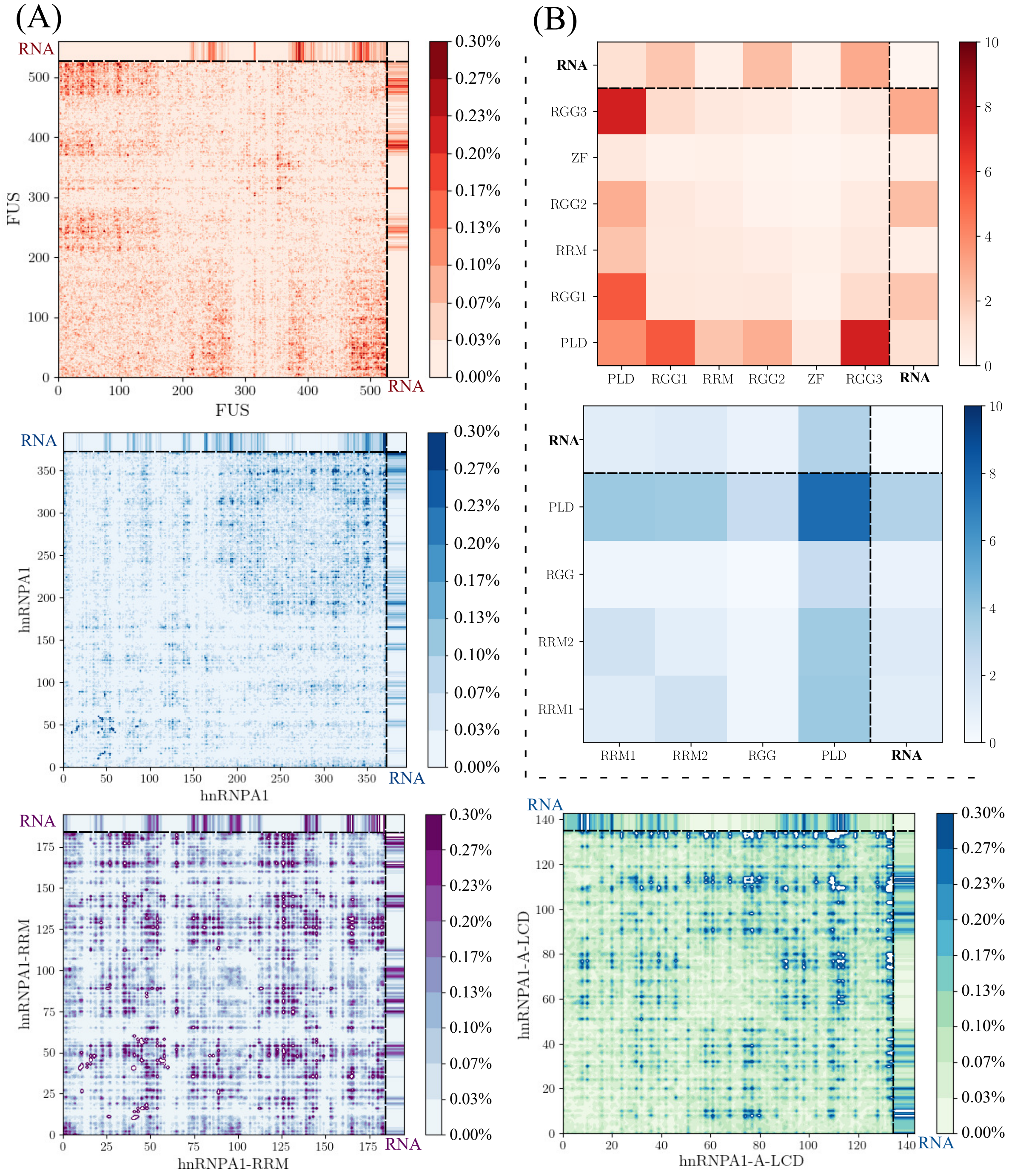


**FIG. S12:** (A) Average number of intermolecular contacts per protein in percentage (where 100% would mean that all protein replicas have a given contact in every configuration) for FUS, hnRNPA1, hnRNPA1-A-LCD and hnRNPA1-RRM condensates in presence of poly-U(250) at $T/T_c = 0.9$ and at the coexisting droplet equilibrium density at such $T$ (see Table S3 for the corresponding critical temperatures of each system). The number of poly-U(250nt) chains are 2 in all systems except for hnRNPA1-RRM where is 1 poly-U(250nt) strand. The corresponding poly-U/protein mass fractions of each system are the following: FUS (0.119), hnRNPA1 (0.099), hnRNPA1-A-LCD (0.117) and hnRNPA1-RRM (0.046). The contacts with poly-U are also included in the upper and right side edges of the maps. (B) Contact map per protein domains for FUS and hnRNPA1 condensates in presence of poly-U at the same conditions described in (A) The protein contact domains with poly-U are also given in the upper and right side edges of the maps.

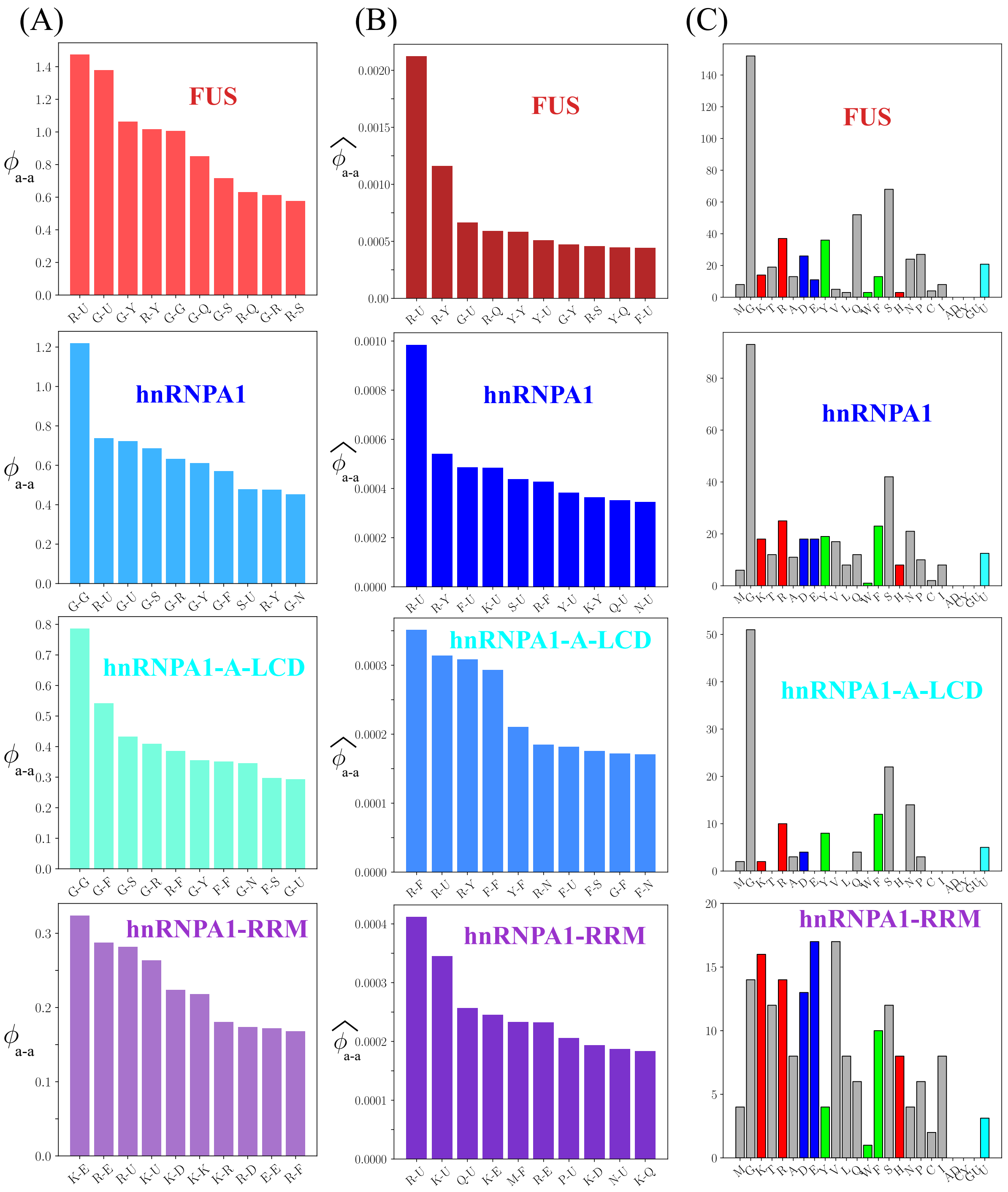


**FIG. S13:** (A) Average number of the ten most repeated intermolecular contacts per protein ($\phi_{a-a}$) in FUS, hnRNPA1, hnRNPA1-A-LCD and hnRNPA1-RRM poly-U condensates at the same concentration and conditions described in Fig. S12. (B) The same as in (A) but renormalized by the protein amino acid and poly-U abundance ($\hat{\phi_{a-a}}$). Note that for every given pairwise contact interaction, we renormalize by the sum of the amount of the two residues/nucleotides involved in the contact interaction divided by two. If the amino acid pair is between the same type of amino acids, we then recover the natural abundance of that amino acid type in the sequence. (C) Abundance of each amino acid (and nucleotide) per protein replica in the system. Red bars indicate the positively charged amino acids, blue color depict negatively charged amino acids, green accounts for aromatic residues and grey for the rest of amino acids. The number of uridines per protein is indicated in cyan.

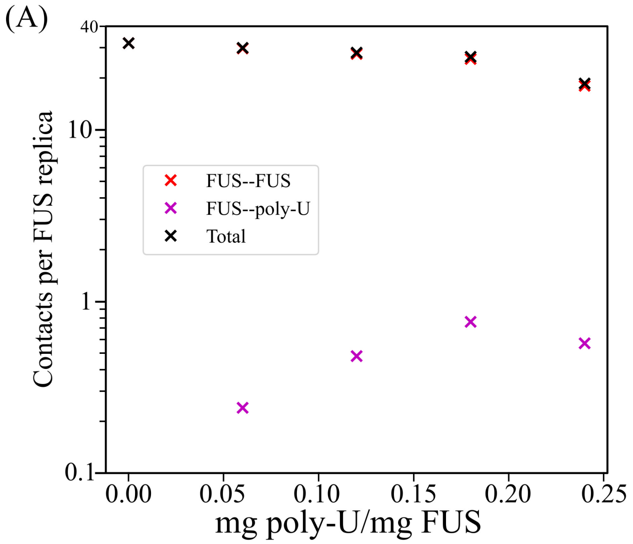


**FIG. S14:** (A) Number of contacts per FUS protein measured in the condensate at $T/T_c = 0.97$ for different concentrations of poly-U of 125 nt length. We include FUS–FUS, FUS–poly-U and the total number of contacts. Poly-U–poly-U contacts have been omitted since they are close to zero due to repulsive electrostatic interactions. (B) Number of contacts measured in the condensate at $T/T_c = 0.97$ for different lengths of poly-U strands and for a constant poly-U/FUS mass ratio of $\sim 0.06$. We include FUS–FUS, FUS–poly-U and the total number of contacts. The limit of the RNA critical length adopted from the obtained critical temperatures in Fig. 4 of the main manuscript is depicted by a dashed vertical line.

### SVII. CALCULATION OF TRANSPORT PROPERTIES WITHIN THE CONDENSATES

We evaluate droplet viscosity and protein diffusion inside the condensates just below the pure component critical temperature from absence to moderately high poly-U concentration for FUS, hnRNPA1, and hnRNPA1-A-LCD proteins. Also, we characterize the viscoelastic properties of wt-TDP-43 condensates without poly-U. We perform NVT simulations in a cubic box at the equilibrium bulk droplet density corresponding to the temperature and poly-U concentration of each system, taken from the phase diagram. We prepare the initial configuration of these systems by compressing the long side of the DC simulation box until obtaining a cubic box, and then the system is relaxed in the NpT ensemble until the bulk equilibrium droplet density is reached. Then, systems are further equilibrated for ∼100 nanoseconds now in the NVT ensemble and, finally, production runs entail from 3 to 6 microseconds depending on the system (around 6 microseconds were simulated for each poly-U/FUS mixture, and almost 5 for each poly-U/hnRNPA1 and poly-U/hnRNPA1-A-

LCD condensates).

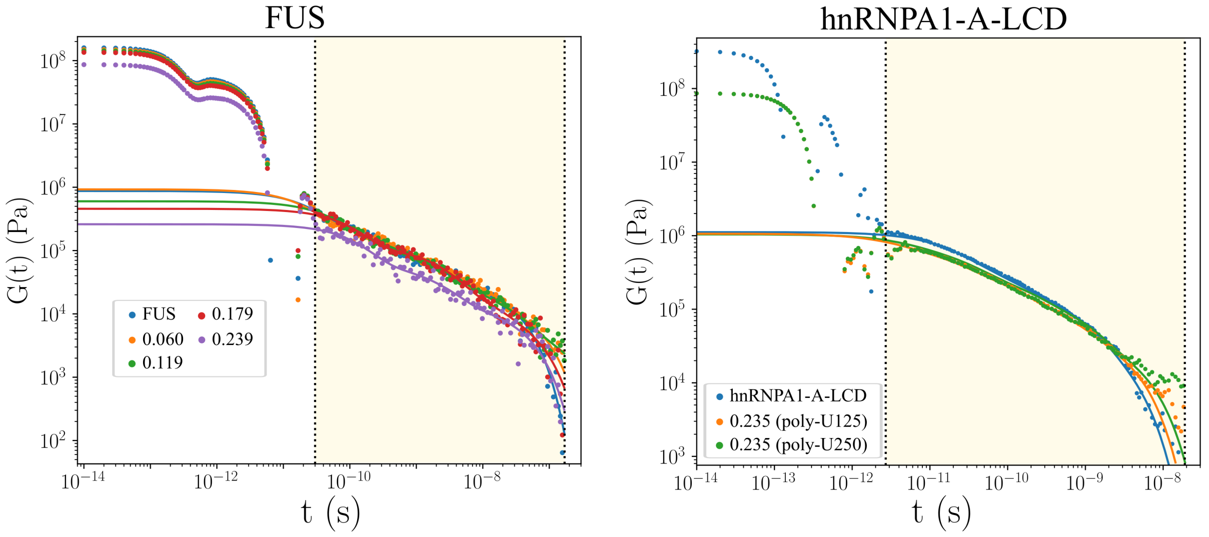


**FIG. S15:** Shear stress relaxation modulus of FUS and hnRNPA1-A-LCD droplets at different poly-U/protein mass fractions as indicated in the legend. Circles show the computed data from NVT simulations at $T/T_c = 0.97$ for both proteins (where $T_c$ refers to the critical temperature of the pure condensates of each protein type) and at the corresponding equilibrium density of each droplet at such conditions. The shadowed regime (light yellow) indicates the region in which the Maxwell modes fit is applied (solid lines). Note that these fits are only applied to compute the contribution to the total viscosity along the shadowed regime. While for hnRNPA1-A-LCD, we plot $G(t)$ for a given RNA concentration using strands of 125 and 250 nt, as well as for the pure protein system, for FUS, all the different concentrations were achieved by adding 125 nt poly-U strands.

From NVT simulations, we can compute both viscosity and protein diffusion in the condensate in separate ways. The shear viscosity can be straightforwardly calculated by integrating the relaxation modulus in time (see Chapter 7 of the book [18]):

$$\eta = \int_0^\infty dt\, G(t) \tag{S10}$$

In an isotropic system, we can compute the shear relaxation modulus $G(t)$ more accurately by using all the components of the pressure tensor ($\sigma_{\alpha\beta}$) as shown in Ref. [19]:

$$\begin{aligned} G(t) &= \frac{V}{5k_BT}\left[\langle\sigma_{xy}(0)\sigma_{xy}(t)\rangle + \langle\sigma_{xz}(0)\sigma_{xz}(t)\rangle + \langle\sigma_{yz}(0)\sigma_{yz}(t)\rangle\right] \\ &+ \frac{V}{30k_BT}\left[\langle N_{xy}(0)N_{xy}(t)\rangle + \langle N_{xz}(0)N_{xz}(t)\rangle + \langle N_{yz}(0)N_{yz}(t)\rangle\right], \end{aligned} \tag{S11}$$

where $N_{\alpha\beta} = \sigma_{\alpha\alpha} - \sigma_{\beta\beta}$ is the first normal stress difference. This correlation can be easily computed by using the compute ave/correlate/long in the USER-MISC package of LAMMPS [1]. In all cases, the relaxation modulus presents an initial regime that mainly accounts for the protein/poly-U intramolecular interactions, followed by a terminal region which corresponds to much slower relaxation modes as those coming from intermolecular interactions and the relaxation of the protein and RNA conformations. Due to the very wide range time-scale involved in the calculation and the noisy nature of the relaxation modulus in the terminal region obtained in the simulations, we follow a particular strategy to calculate our estimate of viscosity. At short times, $G(t)$ is smooth and the integral can be computed using numerical integration (trapezoidal rule). However, at longer times $G(t)$ presents more noise, and hence, we calculate the integral in that regime by first fitting $G(t)$ to a series of Maxwell modes ($G_i \exp(-t/\tau)$) equidistant in logarithmic time [20] and then calculating the integral analytically. The fit to Maxwell modes is carried out with the help of the open-source RepTate software [21]. Finally, viscosity is obtained by adding the two terms:

$$\eta = \eta(t_0) + \int_{t_0}^{\infty} dt\, G_M(t), \tag{S12}$$

where $\eta(t_0)$ corresponds to the computed term for short time-scales, $G_M(t)$ is the part evaluated via the Maxwell mode fit at long time-scales, and $t_0$ is the time that separates both regimes. In Fig. S15 we plot the stress relaxation function $G(t)$ for FUS and hnRNPA1-A-LCD for different poly-U concentrations and lengths. We only fit $G(t)$ in the shadowed regime, and then we integrate it as explained in Eq. (S12). The time $t_0$ is defined by the left dotted vertical line. Note that, due to the finite size of the simulation box and the finite length of the run, the relaxation modulus $G(t)$ shown in Fig. S15 is noisy in the region closer to the terminal time (the time where the modulus decays exponentially to zero). The reported value of the viscosity is the result of the Maxwell mode fit to the noisy $G(t)$ and, thus, has some level of uncertainty. The error bars shown in Fig. 5 of the main paper have been estimated from the error of the Maxwell mode fits to the value of $G(t)$ obtained in simulations.

In Tables S4 and S5 we provide the values of the viscosity measured as a result of the contribution at short times and the integral of the Maxwell mode fit. This results are plotted in Fig. 5 of the main text, but here we also include all values for poly-U strands of 250 nt.

| poly-U / Protein | No RNA | 1x250 (nt) | 2x250 (nt) | 3x250 (nt) | 4x250 (nt) |
|---|---|---|---|---|---|
| FUS | $1.27 \times 10^{-3}$ | $1.80 \times 10^{-3}$ | $2.30 \times 10^{-3}$ | $2.60 \times 10^{-3}$ | $2.30 \times 10^{-3}$ |
| hnRNPA1 | $7.5 \times 10^{-4}$ | $7.0 \times 10^{-4}$ | $6.8 \times 10^{-4}$ | $8.9 \times 10^{-4}$ | $8.9 \times 10^{-4}$ |
| hnRNPA1-A-LCD | $2.92 \times 10^{-4}$ | $3.12 \times 10^{-4}$ | $3.50 \times 10^{-4}$ | $2.90 \times 10^{-4}$ | $3.16 \times 10^{-4}$ |

**TABLE S4:** Viscosity ($Pa \cdot s$) of FUS (at $T/T_c = 0.97$), hnRNPA1 (at $T/T_c = 0.985$) and hnRNPA1-A-LCD (at $T/T_c = 0.98$) condensates as a function of poly-U concentration, given in number of added poly-U strands of 250 nucleotides. The number of proteins in each system is that given in Table S1, and is constant for all concentrations. The equivalence in poly-U/protein mass ratio can be extracted from Fig. S4

| poly-U / Protein | 2x125 (nt) | 4x125 (nt) | 6x125 (nt) | 8x125 (nt) | 26x10 (nt) |
|---|---|---|---|---|---|
| FUS | $1.62 \times 10^{-3}$ | $1.92 \times 10^{-3}$ | $1.22 \times 10^{-3}$ | $0.97 \times 10^{-3}$ | $0.07 \times 10^{-3}$ |
| hnRNPA1 | $5.5 \times 10^{-4}$ | $5.9 \times 10^{-4}$ | $5.4 \times 10^{-4}$ | $4.3 \times 10^{-4}$ | - |
| hnRNPA1-A-LCD | $3.33 \times 10^{-4}$ | $3.00 \times 10^{-3}$ | $2.90 \times 10^{-3}$ | $2.68 \times 10^{-3}$ | - |

**TABLE S5:** Viscosity ($Pa \cdot s$) of FUS (at $T/T_c = 0.97$), hnRNPA1 (at $T/T_c = 0.985$) and hnRNPA1-A-LCD (at $T/T_c = 0.98$) condensates as a function of poly-U concentration, given in number of poly-U strands of 125 nucleotides and 10 nucleotides for the last column. The number of proteins in each system is that given in Table S1, and is constant for all concentrations. The equivalence in poly-U/protein mass ratio is provided in Fig. S4

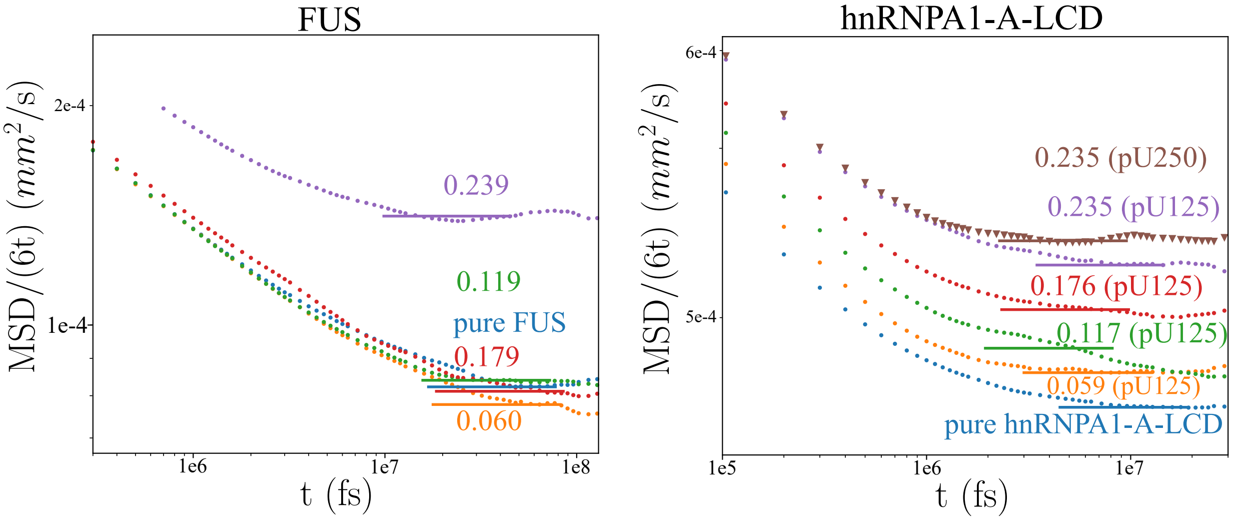


**FIG. S16:** Diffusion coefficients (MSD/6t) of FUS and hnRNPA1-A-LCD proteins inside the condensates at different poly-U/protein mass fractions as indicated in the different curves. The data shown for FUS correspond to poly-U strands of 125 nucleotides length, while for hnRNPA1-A-LCD condensates, strands of 125 and 250 nt were introduced as specified in each curve label. Simulations were performed in the NVT ensemble at $T/T_c = 0.97$ for FUS, and at $T/T_c = 0.98$ for hnRNPA1-A-LCD proteins (where $T_c$ refers the critical temperature of pure condensates) and at the corresponding equilibrium density of each droplet at such conditions. The horizontal solid lines depict where the diffusive regime starts, and thus, where diffusion coefficients can be measured.

The protein diffusion coefficient inside the condensates is obtained through the mean squared displacement (MSD) of the proteins center of mass. After the ballistic regime (i.e., $\sim$ 1 molecular diameter), proteins exhibit a diffusive behavior and then the MSD of the center of mass can be measured via:

$$\left\langle (\mathbf{R}_{\rm CM}(t) - \mathbf{R}_{\rm CM}(0))^2 \right\rangle = 6D_c t, \tag{S13}$$

where $R_{CM}$ indicates the center of mass of a given protein at different times, and $D_c$ accounts for the diffusion coefficient. In order to get an accurate estimate of the MSD, the same correlator technique employed in the calculation of the relaxation modulus has been used [19]. By plotting the MSD divided by $6t$ ( as shown in Fig. S16), the function shows a plateau at long times that provides the value of the diffusion coefficient of the proteins ($D_c$). In contrast to viscosity, protein diffusion coefficients barely depend on the length of the added poly-U strands, but depend on the induced droplet density by each poly-U concentration.

---

[10] J. S. Rowlinson and B. Widom, *Molecular theory of capillarity* (Courier Corporation, 2013).

[11] W. Humphrey, A. Dalke, and K. Schulten, Vmd: visual molecular dynamics, Journal of molecular graphics **14**, 33 (1996).

[12] A. Ladd and L. Woodcock, Triple-point coexistence properties of the lennard-jones system,

Chemical Physics Letters **51**, 155 (1977).

[13] R. García Fernández, J. L. Abascal, and C. Vega, The melting point of ice i h for common water models calculated from direct coexistence of the solid-liquid interface, The Journal of chemical physics **124**, 144506 (2006).

[14] F. J. Blas, L. G. MacDowell, E. de Miguel, and G. Jackson, Vapor-liquid interfacial properties of fully flexible lennard-jones chains, The Journal of chemical physics **129**, 144703 (2008).

[15] J. R. Espinosa, E. Sanz, C. Valeriani, and C. Vega, On fluid-solid direct coexistence simulations: The pseudo-hard sphere model, The Journal of chemical physics **139**, 144502 (2013).

[16] J. Walton, D. Tildesley, J. Rowlinson, and J. Henderson, The pressure tensor at the planar surface of a liquid, Molecular physics **48**, 1357 (1983).

[17] J. M. Prausnitz, R. N. Lichtenthaler, and E. G. De Azevedo, *Molecular thermodynamics of fluid-phase equilibria* (Pearson Education, 1998).

[18] M. Rubinstein, R. H. Colby, *et al.*, *Polymer physics*, Vol. 23 (Oxford university press New York, 2003).

[19] J. Ramírez, S. K. Sukumaran, B. Vorselaars, and A. E. Likhtman, Efficient on the fly calculation of time correlation functions in computer simulations, The Journal of chemical physics **133**, 154103 (2010).

[20] A. E. Likhtman, Single-Chain Slip-Link Model of Entangled Polymers:˜Simultaneous Description of Neutron Spin-Echo, Rheology, and Diffusion, Macromolecules **38**, 6128 (2005).

[21] V. A. Boudara, D. J. Read, and J. Ramírez, Reptate rheology software: Toolkit for the analysis of theories and experiments, Journal of Rheology **64**, 709 (2020).
